# Modeling Avian Full Annual Cycle Distribution and Population Trends with Citizen Science Data

**DOI:** 10.1101/251868

**Authors:** Daniel Fink, Tom Auer, Alison Johnston, Viviana Ruiz-Gutierrez, Wesley M. Hochachka, Steve Kelling

**Affiliations:** Cornell Lab of Ornithology, Cornell University, Ithaca, New York, USA

**Keywords:** biodiversity monitoring, abundance, full annual cycle, bird distributions, population trends, area of occurrence, Wood Thrush, bird migration, eBird, citizen science

## Abstract

Information on species’ distributions and abundances, and how these change over time are central to the study of the ecology and conservation of animal populations. This information is challenging to obtain at relevant scales across range-wide extents for two main reasons. First, local and regional processes that affect populations vary throughout the year and across species’ ranges, requiring fine-scale, year-round information across broad — sometimes hemispheric — spatial extents. Second, while citizen science projects can collect data at these scales, using these data requires appropriate analysis to address known sources of bias. Here we present an analytical framework to address these challenges and generate year-round, range-wide distributional information using citizen science data. To illustrate this approach, we apply the framework to Wood Thrush (*Hylocichla mustelina*), a long-distance Neotropical migrant and species of conservation concern, using data from the citizen science project eBird. We estimate occurrence and relative abundance with enough spatiotemporal resolution to support inference across a range of spatial scales throughout the annual cycle. Additionally, we generate intra-annual estimates of the range, intra-annual estimates of the associations between species and the local environment, and inter-annual trends in relative abundance. This is the first example of an analysis to capture intra- and inter-annual distributional dynamics across the entire range of a broadly distributed, highly mobile species.

## (a) Introduction

Information on the factors that determine species’ distribution and abundance constitutes the foundation of much of our ecological knowledge on animal populations. To date, much of this information has been largely restricted to static patterns across large-spatial extents (e.g. distributions during the breeding season), or dynamic patterns for small-spatial scales (e.g. local extinction and colonization dynamics). However, environmental factors that drive population dynamics are known to vary regionally and seasonally, and failing to account for this variation might yield biased information needed to inform future biodiversity scenarios under changing environmental and climatic conditions. To rise to this challenge, we need to develop analytical frameworks to generate accurate information on species distribution and abundance at relevant spatiotemporal *scales*, i.e. scales at which environmental processes operate, and across the broad spatiotemporal *extents* over which these processes vary (Heffernan *et al.* 2014), i.e. across the entire distributional range of species.

Our ability to model abundance and distribution at relevant spatiotemporal resolutions and extents is largely limited by data availability for many taxonomical groups (Chandler *et al.* 2017, Hortal *et al.* 2015). This is largely driven by the lack of sufficient quantities of high-resolution observational data across broad spatiotemporal extents. Moreover, current information on species distributions often suffer from strong regional biases in data collection (Boakes *et al.* 2010). As a result, information on species distributions is not well suited to capture population dynamics between seasons or between years. Even for birds, one of the best-surveyed taxonomical groups, the majority of our knowledge is restricted to those time periods of data availability (e.g. the breeding season; Marra *et al.* 2015) and we often struggle to track rapidly changing distributions (Massimino *et al.* 2015). There are a few notable exceptions of large-scale monitoring programs that are able to generate continental-scale trends in abundance and distributions (North America: Sauer & Link 2011; Europe: European Bird Census Council 2016). However, these are still restricted to only one stage of the annual life cycle, and these sampling schemes does not go far beyond existing political boundaries to cover the entire distributional range of most species of interest.

Citizen science projects that use crowdsourcing techniques to engage the public are an increasingly reliable source of much needed information for modeling the dynamics of species abundance and distribution, for they have been very successful at collecting observational data across large areas and throughout all seasons (Dickinson *et al.* 2010). However, using these data to generate robust distributional information (at any spatiotemporal scale) is fraught with analytical challenges (Hochachka *et al.* 2012; Bird *et al.* 2014). These challenges have led to the development and application of a number of analytical approaches. For example, to deal with heterogeneous and imperfect observation processes, authors include relevant fixed and random effects (Sauer & Link, 2011; Johnston *et al.* 2018) or rely on explicit models of the detection process (Kéry & Royle, 2015). When project participants choose where and when to conduct their surveys, site selection biases lead to repeated surveys in popular locations and few surveys in areas that are hard to access, contain low species numbers, or few species of interest (e.g, urban centers, grazing or agricultural lands). However, recent data-sampling methods have proven useful to mitigate the effects of these site-selection biases (Robinson *et al.* 2017; Johnston *et al.* 2019). Using citizen science data to estimate abundance also presents the statistical challenge of “zero inflation” where many zero counts can degrade model performance. A wide variety of new abundance models have been proposed to deal with zero-inflation (Denes *et. al.* 2015).

The majority of the methodological developments discussed so far have been used to study regional-scale patterns of species’ distribution and abundance during a single season of the year. Generating distributional information across larger spatiotemporal extents with citizen science data presents three additional challenges: 1) the need to consider large sets of potential environmental factors across a species’ distributional range and annual life cycle; 2) strong variation in data density across large regions; and 3) the spatial and temporal variation in a species’ response to the same environmental conditions. Machine and statistical learning models have proven to be efficient tools for addressing the first challenge, and have proven to be successful at learning complex species-environment relationships from large sets of environmental covariates (Elith & Leathwich 2009). Adaptive knot designs (Gelfand *et al.* 2012) and partitioning methods (Fink *et al.* 2013) have been proposed to deal with the second challenge of variation in data density, which can degrade model performance, by adding multi-scale structure to broad extent analyses. Lastly, non-stationary regression techniques function to add multi-scale structure to analyses and have proven to be a useful solution to the third challenge of variation in response, which can also degrade model performance (Fink *et al.* 2010; Finley 2011).

While previous studies have dealt with one or two of the analytical challenges outlined above (e.g. Johnston *et al.* 2015), none have dealt with all of these challenges simultaneously at the relevant scales necessary to make accurate inferences on species abundance and distributions for broadly distributed species across the full annual cycle. Here, we describe an analytical framework capable of estimating species’ occurrence and relative abundance, year-round and range-wide with enough spatial and temporal resolution to support inference across a range of scales. This includes seasonal predictions of distributions (quantified as the area of occurrence), seasonal estimates of the associations between species and features of their local environment, and year-to-year trends in relative abundance.

As a case study, we analyzed data from the global citizen science project eBird (Sullivan *et al.* 2014) for the long-distance migratory songbird, Wood Thrush (*Hylocichla mustelina*) that breeds in eastern North America and winters largely in Mesoamerica. The Wood Thrush is a species of conservation concern, having suffered steep population declines over the past several decades (Sauer *et al.* 2017). Despite numerous regional studies (e.g. Rushing *et al.* 2017), comprehensive information on patterns of relative abundance, distribution and trends are lacking and much needed to provide a unifying framework for other sources of information (e.g. movement and connectivity). Although we present a case study focused on a bird species, the exponential growth of available citizen science data on other taxonomical groups will render our proposed framework highly useful in future studies of other species and taxonomical groups.

## (b) Materials and methods

### Observational Data

The bird observation data were obtained from the global bird monitoring project, eBird (Sullivan *et al.* 2014) using the eBird Reference Dataset (ERD2016, Fink *et al.* 2017). We used a subset of data in which the time, date, and location of the survey period were reported and observers recorded the number of individuals of all bird species detected and identified during the survey period, resulting in a ‘complete checklist’ of species on the survey (Sullivan *et al.* 2009). The checklists used here were restricted to those collected with the ‘stationary’, ‘traveling’, or ‘area search’ protocols from January 1, 2004 to December 31, 2016 within the spatial extent between 180° to 30° W longitude and north of 0° latitude. Area surveys were restricted to those covering less than 56 km^2^. and traveling surveys were restricted to those ≤ 15km. This resulted in a dataset consisting of 11.7 million checklists, of which a random 10% were withheld for model validation (Appendix S1Figure S1).

### Predictor Data

We incorporated three classes of predictors in the models: (1) Five observation-effort predictors to account for variation in detection rates, (2) Three predictors to account for trends at different temporal scales, and (3) 79 environmental predictors from remote sensing data to capture associations of birds with elevation and a variety of habitats across the continent. The effort predictors were: (a) the duration spent searching for birds, (b) whether the observer was stationary or traveling, (c) the distance traveled during the search, (d) the number of people in the search party, and (e) the checklist calibration index, a standardized measure indexing differences in behavior among observers (Kelling et al. 2015; Johnston et al. 2018). The observation time of the day was used to model variation in availability for detection, e.g. variation in behavior such as participation in the dawn chorus (Diefenbach *et al.* 2007). The day of the year (1-366) on which the search was conducted was used to capture intra-annual variation and the year of the observation was included to account for inter-annual variation.

The environmental predictors included variables describing elevation, topography and land cover. To account for the effects of elevation and topography, each checklist location was associated with elevation, eastness, and northness. These latter two topographic variables combine slope and aspect to provide a continuous measure describing geographic orientation in combination with slope at 1km^2^ resolution (Amatulli *et al.* 2017). Each checklist was also linked to a series of covariates derived from the NASA MODIS land cover data (Friedl *et al.* 2010). We selected this data product for its moderately high spatial resolution, annual temporal resolution, and global coverage. We used the University of Maryland (UMD) land cover classifications (Hansen *et al.* 2000) and derived water cover classes from the MODIS Land Cover Type QA Science Dataset resulting in a class label for each 500m pixel into one of 19 classes (Table 1).

**Table 1:** Land and water cover classes used for distribution modeling. All cover classes were summarized within a 2.8km × 2.8km (784 hectares) landscape centered on each checklist location. Within each landscape, we computed the proportion of each class, and three descriptions of the spatial configuration of the class within the landscape.

|  | Land Cover | Water Cover |
| --- | --- | --- |
| 1 | Evergreen Needleleaf Forest | Shallow Ocean |
| 2 | Evergreen Broadleaf Forest | Ocean coastlines and lake shores |
| 3 | Deciduous Needleleaf Forest | Shallow inland water |
| 4 | Deciduous Broadleaf Forest | Deep Inland Water |
| 5 | Mixed Forest | Moderate or continental ocean |
| 6 | Closed Shrublands | Deep Ocean |
| 7 | Open Shrublands |  |
| 8 | Woody Savannas |  |
| 9 | Savannas |  |
| 10 | Grasslands |  |
| 11 | Croplands |  |
| 12 | Urban and built-up |  |
| 13 | Barren |  |

Checklists were linked to the MODIS data by-year from 2004-2013, capturing inter-annual changes in land cover. The checklist data after 2013 were matched to the 2013 data, as MODIS data after 2013 were unavailable at the time of analysis. All cover classes were summarized within a 2.8km × 2.8km (784 hectare) neighborhood centered on the checklist location. In each neighborhood, we computed the proportion of each class in the neighborhood (PLAND). To describe the spatial configuration of each class we computed three further statistics using FRAGSTATS (McGarigal *et al.* 2012; VanDerWal *et al.* 2014): LPI an index of the largest contiguous patch, PD an index of the patch density, and ED an index of the edge density. Together these four metrics for each of 19 land cover categories led to 76 covariates describing the environment within the local neighborhood.

### Analysis Overview

To meet the analytical challenges of modeling with eBird data, we adopted an ensemble modeling strategy based on the Adaptive Spatio-Temporal Exploratory Model (AdaSTEM; Fink *et al.* 2013). AdaSTEM is a framework for analyzing large-scale patterns with an ensemble of regionally and seasonally local regression models. For each of 100 ensemble runs, the data are independently subsampled and the study extent is partitioned using a randomly located and oriented grid. Each grid cell is a spatiotemporal block (or *stixel*) and an independent regression model, called a *base model*, is fit within each stixel.

Ensemble estimates are made by averaging across the corresponding base model estimates. Combining estimates across the ensemble controls for variability between models (Efron 2014), providing a simple way to control for overfitting while naturally adapting to non-stationary relationships between species and their environments (Fink *et al.* 2010). To make ensemble predictions at a particular location and time, predictions are made from the 100 base models, each from a single ensemble partition, and each fit to an independent subsample of local data in space and time. Because data are subsampled for each base model, point-level uncertainty estimates can be produced by examining variation in the suite of base model predictions across the ensemble. All analysis was conducted in R, version 3.4.2 (R Development Core Team 2017) and deployed using Apache Spark 2.1 (Zaharia, *et. al.* 2016).

In the following sections, we describe the AdaSTEM ensemble design, and the spatiotemporal sampling procedure, the base models run within each stixel. Then we describe how we used the ensemble to estimate four population parameters: (1) landscape-scale estimates of occurrence and abundance, (2) landscape-scale estimates of the area of occurrence, (3) regional-scale habitat use and avoidance, and (4) landscape-scale trends in abundance.

### AdaSTEM Ensemble Design

Stixel size controls a bias-variance tradeoff (Fink *et al.* 2013) and must strike a balance between stixels that are large enough to achieve sufficient sample sizes to fit good base models (i.e. limiting variance of estimates), and small enough to assume stationarity (controlling bias). Under the AdaSTEM framework, we set all stixel’s temporal width to 30.5 days. The spatial dimensions were adaptively sized to generate smaller stixels in regions with higher data density, using QuadTrees (Samet 1984), a recursive partitioning algorithm. The AdaSTEM ensemble consisted of 100 randomly located and oriented grids of overlapping spatiotemporal stixels generated in this way. See Appendix S1 in Supporting Information for additional information about the specification of the ensemble design.

### Spatiotemporal Sampling

Within each stixel, a spatial case-control sampling strategy was used to address the challenges of highly imbalanced data and site selection bias. Imbalanced data arise when there are a very small number of species detections and a very large number of non-detections. This is a modeling concern because binary regression methods, like the first component of our ZI-BRT base model, become overwhelmed by the non-detections and perform poorly (King & Zeng 2001, Robinson *et al.* 2017). Case-control sampling treats detection and non-detection cases separately, resampling each case to improve spatial and temporal balance in the data and model performance (e.g. Breslow 1996, Fithian & Hastie 2014). See Appendix S2 for additional information about the spatiotemporal sampling procedure.

For the trend base models, we also balanced the per year sample size, after spatiotemporal case control sampling, to control for potential bias associated with the strong inter-annual increases in eBird data volume, 20-30% per year since 2005. Years with fewer data than the average were over-sampled (i.e. randomly sampled with replacement) and years with more data than the average were under-sampled (i.e. randomly sampled without replacement). This sampling strategy, resulted in a training data set with the same per-year sample size.

### Base Models

Within each stixel, relationships between the species response and the predictor variables were assumed to be stationary. To estimate occurrence and relative abundance from the large predictor set while accounting for high numbers of zero counts, we used a two-step Zero-Inflated Boosted Regression Tree (ZI-BRT) model (Johnston *et. al.* 2015; Ridgeway *et al.* 2017). In the first step a Bernoulli response BRT was trained to predict the probability of occurrence and in the second step a Poisson response BRT was trained to predict expected counts conditional on occurrence. To facilitate the estimation of the binary occurrence state from the predicted occurrence probabilities, we also recorded the threshold that maximized the Kappa statistic (Cohen 1960). All predictors were included in both BRTs. The inclusion of the effort and time covariates allowed the models to account for several important sources of variation in detectability. See Appendix S3 for additional information about base model boosted regression tree parameters.

Base models were trained only when there were at least 50 checklists (prior to oversampling) from the spatially balanced case-control sampling procedure and at least 10 species detections (prior to oversampling). To guard against the effects of replicate surveys at popular birding locations, only one detection per day is considered from each location. Stixels that did not meet these minimum sample size requirements were dropped without replacement from the ensemble leading to fewer overlapping base model estimates, and higher variance among ensemble average estimates in regions with low data density or low species detection rates.

To estimate inter-annual trends in relative abundance we trained a second ZI-BRT base model, identical to the one above except for two important modifications. First, to increase species’ encounter rates and strengthen trend signals we aggregated the training data across a 25.2km x 25.2km grid, separately for each year. The aggregation summed the counts of species seen, the durations spent searching for birds, the distances traveled during the search, the numbers of people in the search party, and the checklist calibration values (weighted by search duration) across all checklists within each grid cell. All other covariates were averaged across all checklists within each grid cell. Second, to control for the inter-annual increases in eBird data volume, the aggregated training data was sampled to have the same number of surveys each year. See the Spatiotemporal Sampling section and Appendix S2 in Supporting Information for more information about the sampling procedure.

### Estimating Occurrence and Relative Abundance

Within each stixel, the binomial BRT submodel was used to predict the expected occurrence rate. The expected relative abundance was estimated as the product of the predicted occurrence and the predicted abundance conditional on occurrence. To control for variation in detection rates, the search effort predictors (search duration, protocol, search length, number of observers, and checklist calibration index) were held constant for the predictions. Additionally, to maximize the species’ availability for detection within each stixel, expected values were calculated for the time of day value which maximized the species’ probability of being reporting, based on the partial dependency estimate for time of day (see Regional Habitat Associations section for information on partial dependence estimates.)

The resulting quantity used to estimate occurrence was defined as the probability that an expert eBird participant (top 1% of checklist calibration indices) would detect the species on a search at the optimal time of day for detection while traveling 1 km on the given day at the given location. Relative abundance was estimated as the expected count of individuals of the species on the same standardized checklist. Although this approach accounts for variation in detection rates, it does not directly estimate the absolute detection probability. For this reason, our estimates of occurrence can only be considered as a *relative* measure of species occupancy. Similarly, we refer to the expected count of individuals of the species on the same standardized checklist as a measure of *relative* abundance. Note that this measure of relative abundance is equivalent in many respects to the relative abundance estimates used to estimate trends with the North American Breeding Bird Survey (Sauer & Link 2011).

The ensemble estimates of occurrence and relative abundance were calculated by averaging across all the base model estimates for a given location and date. We generated two sets of ensemble estimates for relative abundance, one designed for high resolution, year-round population mapping and one designed as the basis of the seasonal trend estimates. For the high resolution, year-round population mapping we estimated occurrence and relative abundance for a single day at the center of each week for all 52 weeks of 2016 for each 2.8km × 2.8km grid cell in the Western Hemisphere. For the seasonal trend estimates, we generated weekly estimates of relative abundance for each week within the specified seasons, separately for each year 2007-2016, within each 25.2km × 25.2km grid cell.

Uncertainty was estimated as the lower 10^th^ and upper 90^th^ percentiles based on the variation in the base model estimates. Ensemble average estimates were not made in areas of low data density, where base model minimum sample size requirements were not met. See Appendix S4 for information about subsampling procedures used to estimate uncertainty of the occurrence and abundance estimates.

### Estimating Area of Occurrence

To estimate the Area of Occurrence (AOO) we tested the binary occupied versus unoccupied state for each week and prediction location using both the 2.8km and 25.2km spatial grids, described above. The resulting set of AOO values provides detailed information about the distributional range of a species and can be used to generate fine-scale range boundaries throughout the year.

At the base model level, each location was considered to be occupied if the predicted occurrence probability was above the kappa-maximized threshold for that base model. Aggregating across the ensemble, a location was considered to be occupied if at least 1/7 of base models predicted it was occupied. This is equivalent to an expert observer detecting the species at least once during 7 adjacent days of standardized surveys, taking account of the variation across base models. See Appendix S5 in Supporting Information for further information about the methods used to estimate AOO.

### Estimating Local Trends

To estimate the average annual rate of change in a species’ relative abundance with moderately high spatial resolution (25.2km × 25.2km) we use a two-step approach that exploits the ensemble structure of AdaSTEM. In the first step, a hypothesis-testing approach uses the variation across the ensemble to filter out regions where the estimated direction of the trend was inconsistent. We call this step the *signal filter*. Then in the areas that passed the signal filter, we averaged across the ensemble to remove the intra-ensemble variation while generating trend estimates.

The signal filter began by generating the base model estimates of the slope of the log-linear regression of relative abundance on year and then testing across the ensemble to determine if the slopes were increasing or decreasing. For those locations where the same direction was consistently observed across the ensemble, we then computed an ensemble averaged estimate of the trend as the percept per year change in population size. This trend was estimated as the slope from the log-linear regression of the ensemble average estimates of relative abundance, as described in the Local Occurrence and Relative Abundance section. See Appendix S6 in Supporting Information for further information about the methods used to estimate local trends.

### Estimating Regional Habitat Associations

For each base model, we quantified the strength and direction of association for each cover class predictor. Predictor importance (PI) statistics measured the strength of the overall contribution of individual predictors as the change in predictive performance between the model that includes all predictors and the same model with permuted values of the given predictor (Breiman 2001). PI statistics capture both positive and negative effects arising from both additive and interacting model components. Partial Dependence (PD) statistics described the functional form of the additive association for each individual cover class predictor by averaging out the effects of all other predictors (Hastie *et al.* 2009). To measure the direction of association, we estimated the slope of each PD estimate using simple linear regression.

To examine how species’ habitat use and avoidance varied among regions and seasons, we computed regional trajectories of the strength and direction of the cover class associations. Given the region and the set of predictors to compare, the PI statistics were standardized to sum to 1 across the predictor set for each base model within the specified region. Then, loess smoothers (Cleveland *et al.* 1992) were used to estimate the trajectories of relative predictor importance throughout the year for each predictor. Similarly, a loess smoother was used to estimate the proportion of increasing PD estimates throughout the year for each predictor. Predictors with proportions of base models greater than 70% were considered to have positive associations with species abundance and predictors with proportions less than 30% were considered to have negative associations with species abundance. Predictors with inconsistent directions, those between 30 and 70%, were excluded from summaries.

To quantify changes in habitat use and avoidance throughout the annual cycle, we made weekly estimates of the association between Wood Thrush occurrence and the amount of each habitat class in the local landscape (Fig 3). For each week, the associations were summarized across the population core area, the 5° longitude × 5° latitude area located at the population center for that week. For each cover class, values were combined for both PLAND and LPI predictors to describe the relative strength and direction of the association. Larger absolute values indicate stronger associations and the sign of the value indicates class use or avoidance. Classes with inconsistent direction of association, were removed, resulting in total weekly relative importance that sums to less than 1.

### Model Validation

To assess the quality of the ensemble estimates of AOO, occurrence, and abundance, we validated the model predictions at 2.8km × 2.8km × 1wk resolution using independent validation data. The statistics were evaluated using a Monte Carlo design of 25 spatially balanced samples to help control for the uneven spatial distribution of the validation data with each week (Fink *et al.* 2010; Roberts *et al.* 2017). To quantify the predictive performance for the AOO we used the Area Under the Curve (AUC) and Kappa (Cohen 1960) statistics to describe the models’ ability to classify occupied versus unoccupied sites (Freeman & Moisen 2008). Thus, these metrics are also useful for assessing the quality of the weekly range boundaries. AUC measures a model’s ability to discriminate between positive and negative observations (Fielding & Bell 1997) as the probability that the model will rank a randomly chosen positive observation higher than a randomly chosen negative one. Cohen’s Kappa statistic (Cohen 1960) was designed to measure classification performance accounting for the background prevalence. To quantify the quality of the occurrence estimate as a rate within areas estimated by the AOO to be occupied, we also evaluated AUC and Kappa. To quantify the quality of the abundance estimates we computed Spearman’s Rank Correlation (SRC) and the percent Poisson Deviance Explained (P-DE). SRC measures how well the abundance estimates rank the observed abundances and the P-DE measures the correspondence between the magnitude of the estimated counts and observed counts.

To validate the methods used to estimate the trends we conducted a simulation analysis to assess performance across a wide range of spatially varying and constant trend scenarios coupled with a realistic data observation process. By comparing estimated trends to the simulated truth, we quantified false detection (type I error) and power (type II error) rates at the 25.2km × 25.2km resolution when identifying locations with increasing and decreasing trends. Additionally, the simulations were designed to assess the ability of the method to identify spatially varying trend patterns. The information generated from the simulation study provides insight about the robustness of the trend analysis. See Appendix S7 for further information about the trend simulation study design.

## (c) Results

### Weekly AOO, Occurrence and Relative Abundance

Using the Wood Thrush as exemplar analysis, we generated estimates of AOO, occurrence and relative abundance at a spatiotemporal resolution of 2.8km × 2.8km × 1 week (Fig. 1). Across the study extent, the AOO shows seasonal changes in the distributional range size and shape while the abundance estimates capture regional and seasonal variation in population structure within the distributional range. The breeding season range fills in the eastern deciduous forests east of the Great Plains with highest population concentrations in the Appalachian Mountains (Fig. 1a). During autumn migration, the population concentrates in the southern part of the Appalachian Mountains (Fig. 1b) before crossing the Gulf of Mexico into Central America. The non-breeding distribution (Fig. 1c) is concentrated in the Yucatán Peninsula, with lower concentrations extending north into Veracruz and south to Costa Rica and Panama. During the spring migration (Fig. 1d), Wood Thrush crosses the Gulf of Mexico, concentrating on the Gulf Coast and again in the southern part of the Appalachian Mountains.

**Figure 1:**
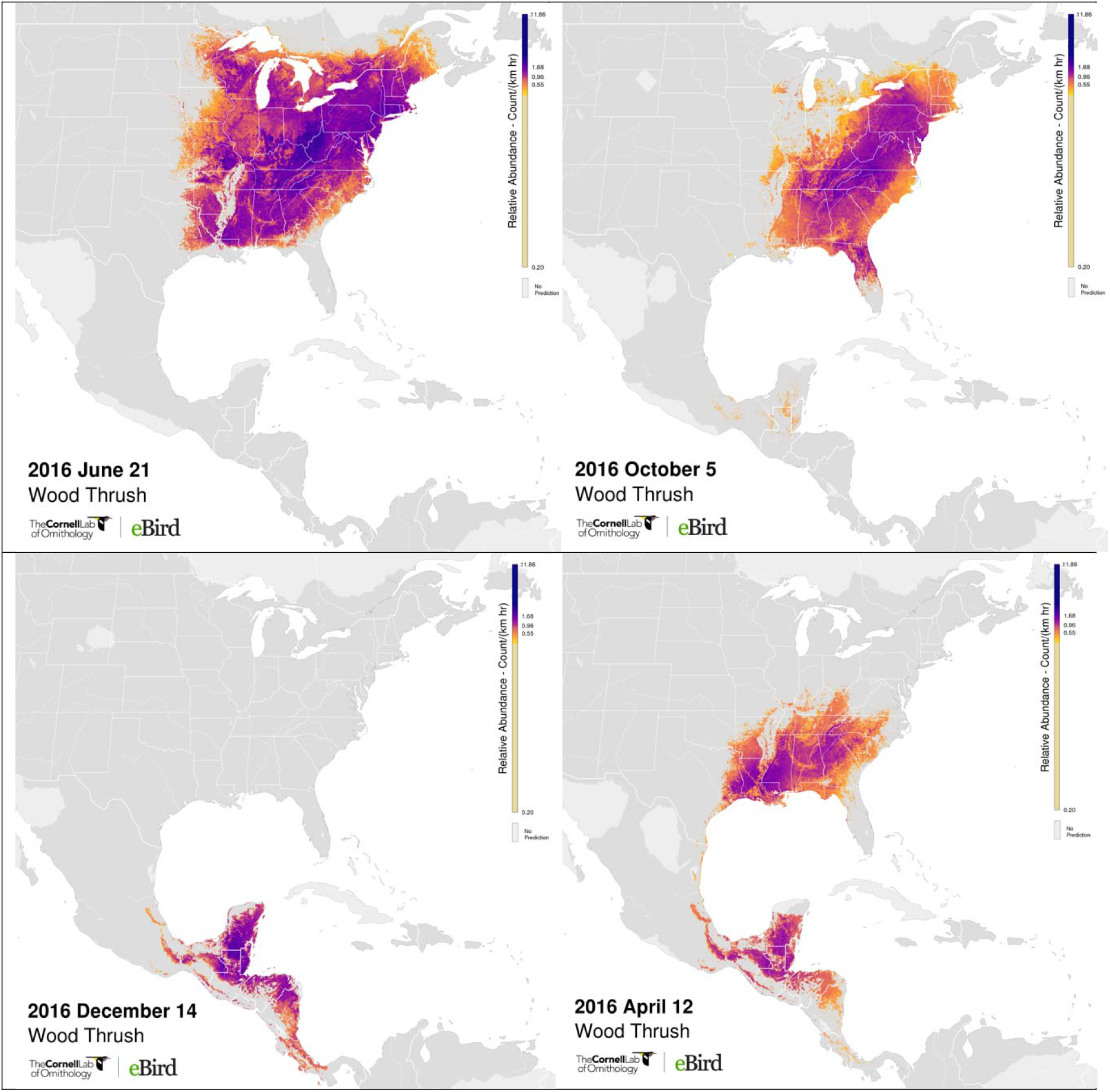
Wood Thrush estimates of Area Of Occurrence (AOO) and relative abundance at 2.8km × 2.8km resolution during (a) breeding (June 20), (b) autumn migration (October 3), (c) non-breeding (December 12), and (d) spring migration (March 28) seasons. Positive abundance is only shown in areas estimated to be occupied and the AOO is depicted as the boundary between pixels with and without color. Brighter colors indicate areas occupied with higher abundance. Relative abundance was measured as the expected count of the species on a standardized 1km survey conducted at the optimal time of day for detection. Note that detectability varies seasonally, complicating comparisons of population size between seasons.

To assess the accuracy of estimates, we calculated range-wide validation estimates based on spatially balanced samples of independent eBird observations for each week of the year. AOO weekly median AUC scores were between 0.73 and 0.91 with mean 0.82 (Fig. 2a) and AOO weekly median Kappa scores were between 0.26 and 0.62 with mean 0.40 (Fig. 2b). Occurrence weekly median AUC scores were between 0.57 and 0.91 with mean 0.72 (Fig. 2c) and occurrence weekly median Kappa scores were between 0 and 0.61 with mean 0.28 (Fig. 2d). Relative abundance weekly median P-DE scores were between 0 and 0.52 with mean 0.19 (Fig. 2e) and relative abundance weekly median SRC scores were between 0.16 and 0.70 with mean 0.41 (Fig. 2f). Weeks with insufficient validation data were shown as zero. These weeks occurred during spring and autumn migration, when detection rates and counts were at their lowest. Variation in predictive performance was highest during the non-breeding season for all metrics, reflecting lower data densities in Mesoamerica.

**Figure 2:**
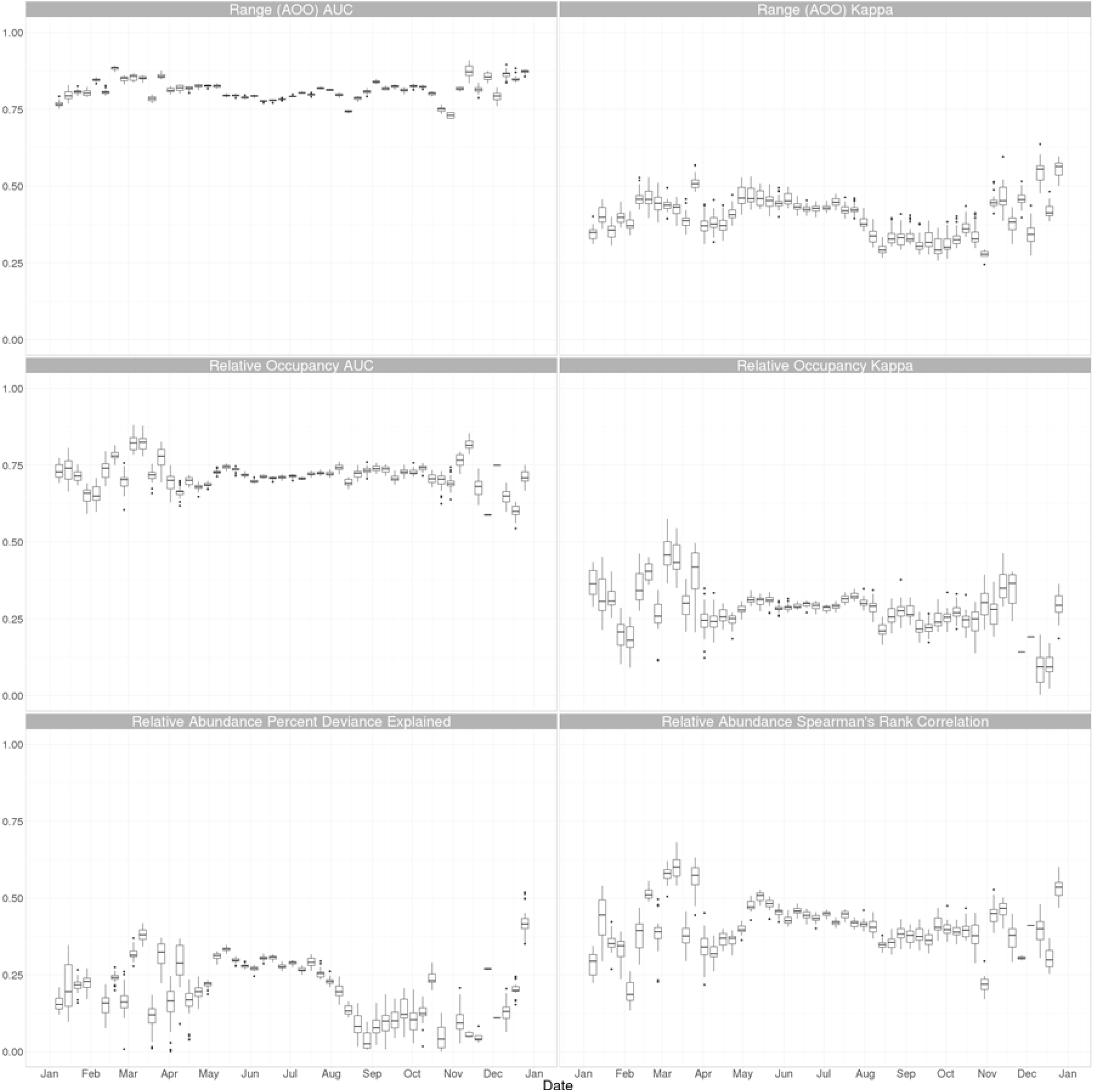
Boxplots of range-wide weekly predictive performance for Area Of Occurrence, occurrence and relative abundance estimates across 25 Monte Carlo samples of spatially balanced validation data. (a) AUC and (b) Kappa scores for area of occurrence estimates. (c) AUC and (d) Kappa scores for occurrence estimates. (e) Spearman’s Rank Correlation and (f) Percent Deviance Explained scores for relative abundance estimates.

### Seasonal Habitat Use and Avoidance

The accuracy of the habitat associations follows from the strong validation results (Fig. 2). The Wood Thrush breeding season is characterized by the strong positive association with deciduous broadleaf forest and the non-breeding season is characterized by the strong positive association with broadleaf evergreen forest (Fig. 3). During spring and autumn migrations, the population is associated with a wider variety of cover classes, and a more even distribution of associations, both positive and negative. This includes a notable positive association with the urban developed class. While these patterns of habitat use and avoidance were consistent with the empirical results documented by Zuckerberg *et al.* (2016) and the qualitative patterns described in Evans *et al.* (2011), they also provided much more detail about the underlying population structure throughout the rest of the year.

**Figure 3:**
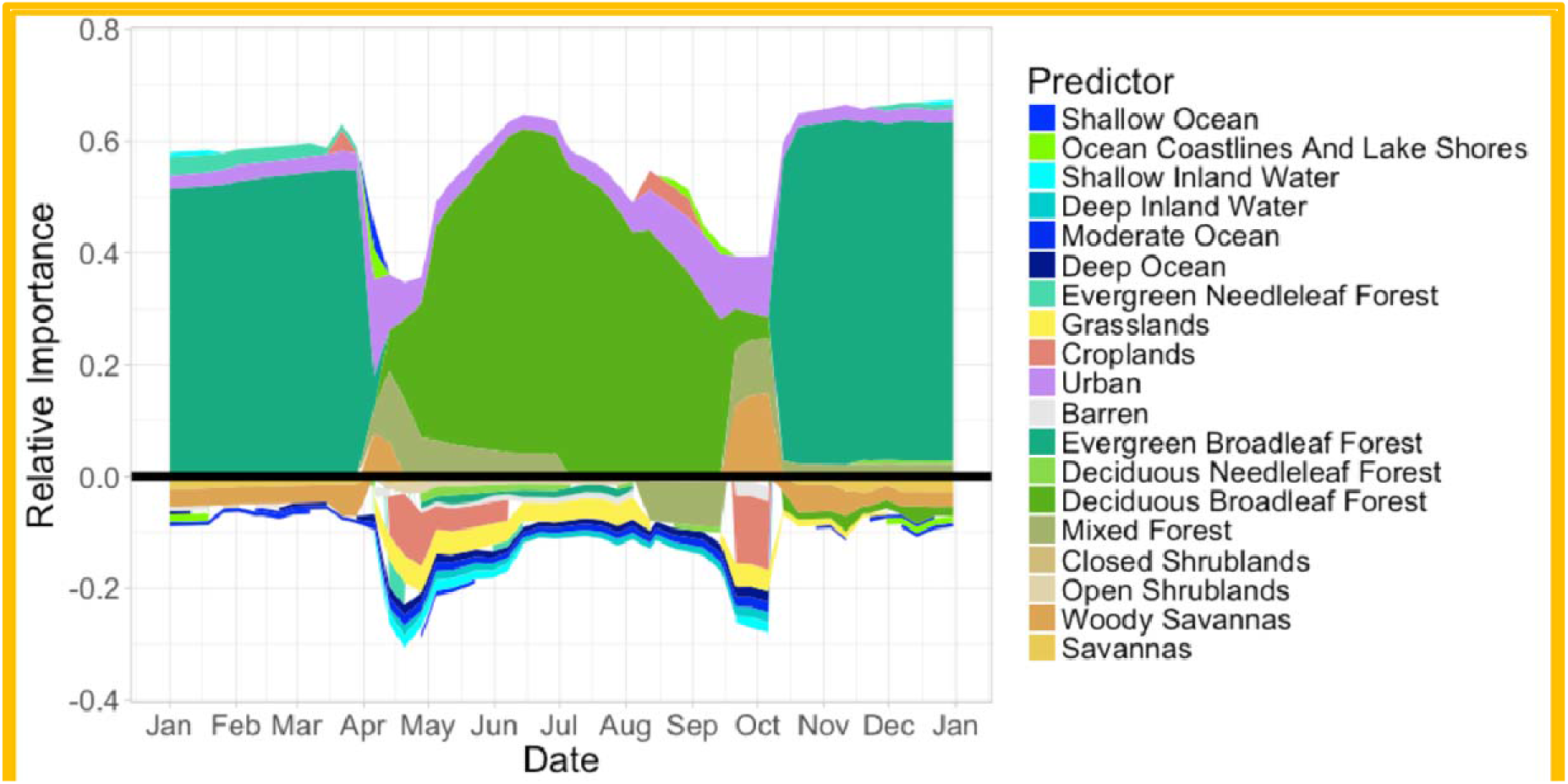
The weekly relative importance for the amount of each land and water cover class for the core Wood Thrush population. Positive importance indicates class use and negative importance indicates class avoidance. The strength of the association with each class is proportional to the width of the class color. Classes with inconsistent direction of association were removed, resulting in total weekly relative importance that sums to less than 1.

### Breeding Season Trends

Fig. 4A shows the average percent change per year in relative abundance from 2007–16 during the breeding season (May 30– July 3). The largest population changes have occurred across the core of the population, the large area of high-abundance including all but the peaks of the Appalachian Mountains (Fig. 4A). Moderate declines of 1 to 3.5% per year were estimated in most locations across this region. However, declines have not occurred range-wide. There were also low abundance portions of the population along the northwest and southern boundaries of the range that increased during the study period. Appendix S6: Fig. S1 shows trend maps of the location-wise upper 2.5% and lower 97.5% confidence limits from the subsampling analysis. These maps show similar moderate-magnitude declines across the core of the breeding range with regional patterns similar to those in Fig. 4A.

**Figure 4:**
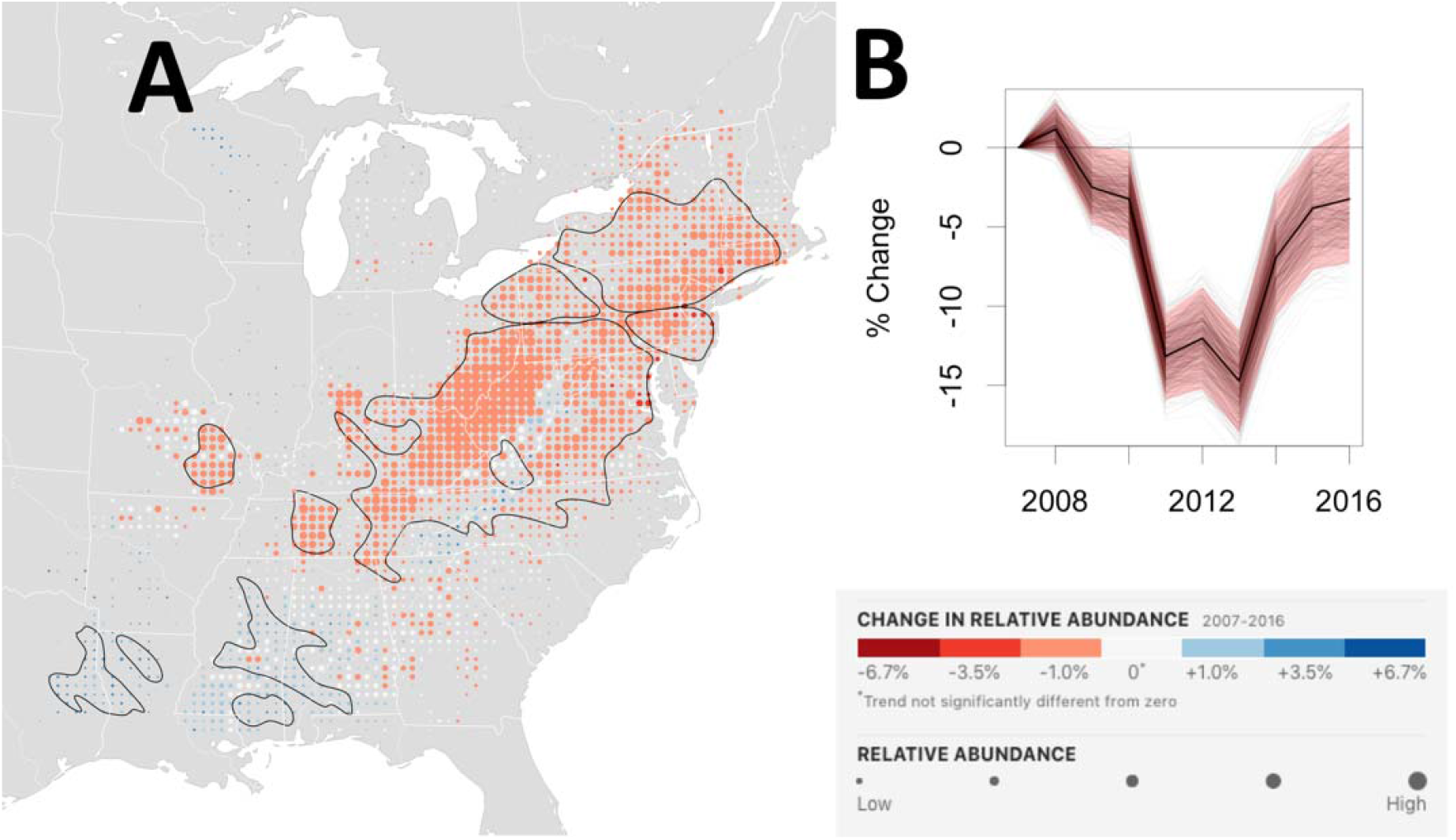
Wood Thrush breeding trend map and range-wide population trajectory. **(A)** The breeding season (May 30– July 3) average annual percentage change in relative abundance from 2007–16. Increases in population size are shown in blue and decreases are shown in red. Darker colors indicate stronger trends. Each dot on the map represents a 25km x 25km area. To help visualize the relative change in population size at each location, the size of each dot has been scaled according to the average abundance at that location during the 10-year study period. Within the regions delineated by the black contour line, the expected False Discovery Rate (type I error) is up to 5% when identifying locations with increasing and decreasing trends. Outside the black contours, the direction of population change is less certain. The breeding season power analysis suggests that regions within the black contours contain 60% of all locations across the breeding range with non-zero trends and contain 80% of all trends with trend magnitudes of 6.7%/*yr* or more (approximately equivalent to halving or doubling of the population across 10 years). **(B)** The trajectory shows the range-wide change in population size starting from 2007. The dark black line is the conditional mean estimate, the red polygon are the 95% confidence limits, and the light grey trajectories show the 500 replicate estimates.

The breeding season range-wide, abundance-weighted trend estimate was −1.48% per year, with a 95% confidence interval between −1.89% and −1.01% per year. The range-wide population trajectory (Fig. 4B) shows the steepest declines in population size between 2010 and 2013 followed by lower rates of decline in the population size from 2013 to 2016.

The simulation study for the breeding season Wood Thrush trends provide information about likely false detection (type I error) and power (type II error) rates when identifying locations with increasing and decreasing trends. The black contour lines in Fig. 4A delineate those regions across which the expected False Discovery Rate is at most 5%. These regions include most of core high-abundance breeding range. The breeding season power analysis (Appendix S7: Fig. S5A) suggests that regions within the black contours contain approximately 60% of all locations across the breeding range with non-zero trends, >67% of trends ≥ |1%/*yr*|, >75% of trends ≥ |3.5%/*yr*|, and 80% of trends ≥ |6.7%/*yr*|. These power results also help with interpretation of results outside the contour lines, providing information about the likely number of locations with trends and the likely strengths of those trends. The breeding season simulation study also suggests that spatially varying trend patterns can be reliably estimated in regions with moderate to strong trends (Appendix S7: Fig. S1 & S2). Overall, these simulation results suggest that there is sufficient data density to estimate moderate to strong trends with low False Discovery Rates (FDR) (Type I errors) and fairly high power (i.e. low Type II errors) across much of the breeding range.

### Nonbreeding Season Trends

Fig. 5A shows the average percent change per year in relative abundance from 2007–16 during the non-breeding season (Dec 1–Feb 28). This map shows moderate declines of 1-3.5% per year across most of the nonbreeding range with the steepest declines in the north eastern part of the Yucatan peninsula and the southern portion of the range extending though eastern Nicaragua, both areas of low abundance. Trend maps of the location-wise upper and lower 95% confidence limits (Appendix S6: Fig. S2) generally show similar spatial patterns with consistent declines surrounding the high abundance population areas centered near the shared boundaries of Mexico, Guatemala, and Belize.

**Figure 5:**
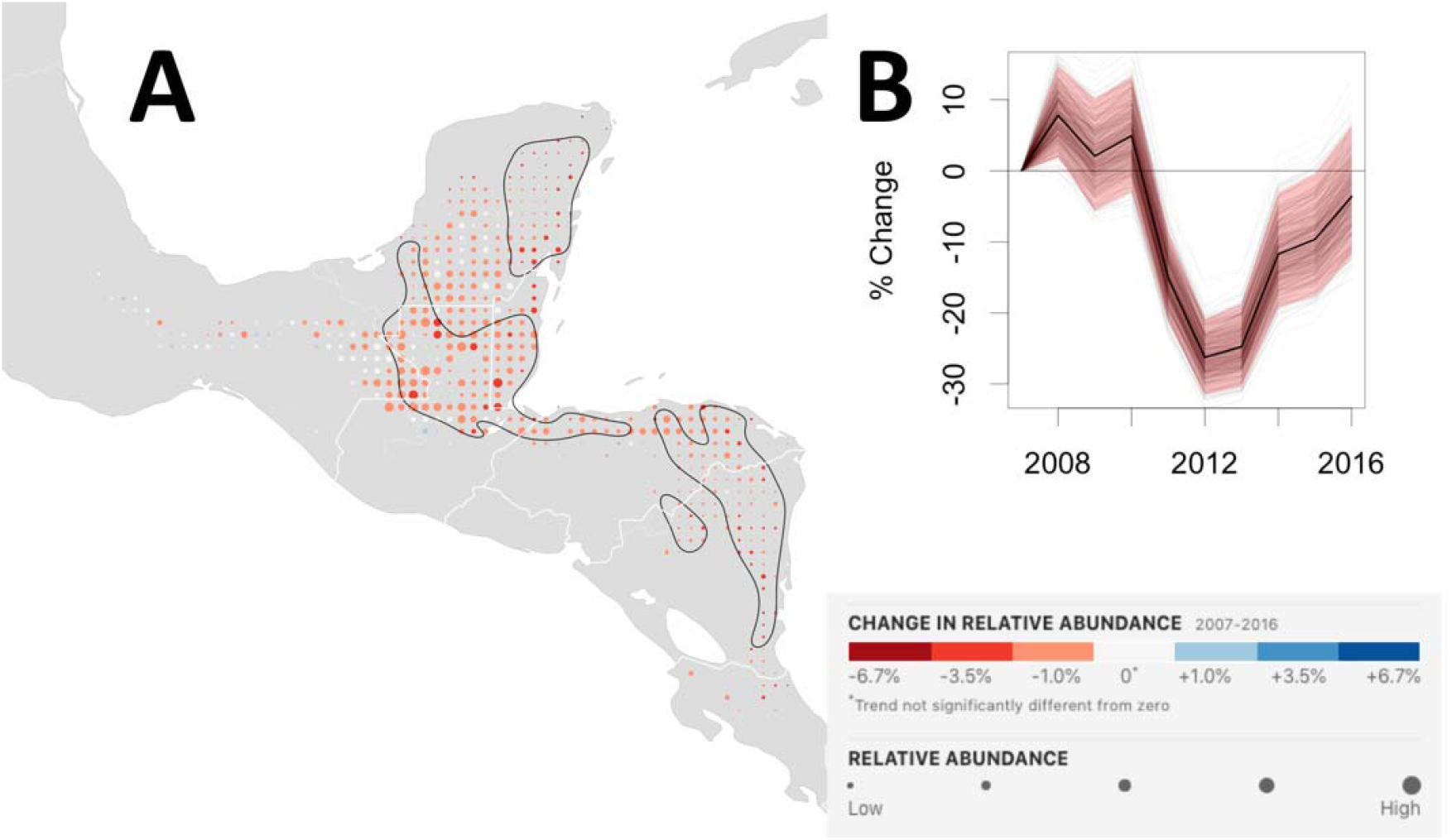
Wood Thrush nonbreeding trend map and range-wide population trajectory. **(A)** The nonbreeding season (Dec 1–Feb 28) average annual percentage change in relative abundance from 2007–16. Increases in population size are shown in blue and decreases are shown in red. Darker colors indicate stronger trends. Each dot on the map represents a 25km x 25km area. To help visualize the relative change in population size at each location, the size of each dot has been scaled according to the average abundance at that location during the 10-year study period. Within the regions delineated by the black contour line, the expected False Discovery Rate (type I error) is up to 5% when identifying locations with increasing and decreasing trends. Outside the black contours, the direction of population change is less certain. The breeding season power analysis suggests that regions within the black contours contain 40% of all locations across the breeding range with non-zero trends and contain 70% of all trends with trend magnitudes of 6.7%/*yr* or more (approximately equivalent to halving or doubling of the population across 10 years). **(B)** The trajectory shows the range-wide change in population size starting from 2007. The dark black line is the conditional mean estimate, the red polygon are the 95% confidence limits, and the light grey trajectories show the 500 replicate estimates.

The range-wide, abundance-weighted nonbreeding season trend estimate qualitatively mirrors the pattern for the breeding season. The estimated nonbreeding season trend is −2.16% per year, with a 95% confidence interval between −2.98% and −1.28% per year. The range-wide population trajectory (Fig. 5B) shows the steepest declines in population size between 2010 and 2013 followed by lower rates of decline in the population size from 2013 to 2016, similar to the range-wide trajectory for the breeding season (Fig. 4B).

The 5% FDR regions delineated by the black contour lines in Fig. 5A surround the high abundance region centered near the shared boundaries of Mexico, Guatemala, and Belize. The nonbreeding season power analysis (Appendix S7: Fig S5B) found that regions within the black contour contain approximately 40% of all locations across the nonbreeding range with non-zero trends, >41% of trends ≥ |1%/*yr*|, 50% of trends ≥ |3.5%/*yr*|, and 70% of trends ≥ |6.7%/*yr*|. These simulation results suggest that there is sufficient data density to estimate strong trends with low FDR and moderate power. The estimated and simulated nonbreeding trend maps presented in Appendix S7: Fig. S3 & S4 suggests that spatially varying trend patterns can be reliably estimated in regions with moderate to strong trends.

## (d) Discussion

In this paper, we show that a combination of semi-structured (Kelling et al. 2019) citizen science data and analyses chosen to deal with the biases in these data can be used to estimate complex patterns of species’ distribution and abundance, at fine spatial and temporal scales, across the full annual cycle. The resolution, extent, and completeness of the information that can be generated with this approach is unprecedented, and has the potential to increase our ecological knowledge and inform conservation plans for a range of species, regions and seasons (Runge *et al.* 2015).

To the best of our knowledge, this is the first comprehensive population-level analysis of distribution, abundance, and habitat use, for Wood Thrush and the first analysis of this kind for Neotropical migrants. The comprehensiveness of the Wood Thrush analysis presented here fills important knowledge gaps, providing novel range-wide and population-level information during the less well-studied migration and the overwintering periods (Evans *et al.* 2011). Moreover, having a comprehensive *quantitative* description of the Wood Thrush population provides the data necessary to track changes in the future and it provides an empirical full-annual-cycle framework with which to integrate other types of data.

We also demonstrate the ability to use citizen science data to estimate trends in relative abundance — a task usually left to monitoring programs which employ more stringent sampling protocols and are hard to deploy across broad extents. The ability to quantify regional differences in trends, especially outside of the breeding season provides important, novel information that can be used to advance our knowledge across multiple fields, from questions about the potential drivers of the evolution of migratory behavior, to providing a framework for the use of information on movement and migratory connectivity to delineate and model contemporary sub-populations.

The year-to-year population trajectories for breeding (Fig. 4B) and nonbreeding (Fig. 5B) seasons follow a very similar pattern, showing the steepest declines in population size between 2010 and 2013 followed by weaker decreases in the population size from 2013 to 2016. The strong correspondence between these independently estimated trajectories provides strong evidence that our results for each life stage are representative of the entire population.

The ability to estimate trends at different times of the year provides essential contextual information to tease apart where in the annual cycle population changes are and are not occurring. Though the average annual rate of population-wide decline is slightly stronger for the nonbreeding (−2.2%) compared to the breeding season (−1.5%), the 95% confidence intervals (−3.0, −1.3%) and (−1.9, −1.0%), respectively, overlap. Given the greater uncertainty and lower power of nonbreeding season analysis results, we do not view this difference as strong evidence of a difference in the rates of decline. This similarity in breeding and non-breeding trends rules out strong inter-annual changes in mortality during the autumn migration.

The nonbreeding trend maps show spatial variation in the pattern of declines, with the steepest declines within the 5% FDR regions (Fig. 5A and Appendix S6: Fig. S2). These results could help tease apart the contrasting results from the demographic models of Taylor & Stutchbury (2016) and the analysis of Rushing *et al.* (2017) on the drivers of population declines of Wood Thrush. Information about regional variation in trends can be used in combination with spatial information on regional threats (e.g. deforestation or other forms of habitat modification) and integrated with information on migratory connectivity to better understand how factors the breeding and non-breeding season influence population declines (e.g. Kramer *et al.* 2018).

More broadly, the analysis presented here demonstrates how citizen science data can be used to generate accurate species-level information for broad-scale biodiversity monitoring like those outlined by the Group on Earth Observations Biodiversity Observation Network (Kissling *et al.* 2017). It is worth noting that without a single, comprehensive source of information, making population-wide assessments requires the additional steps to acquire, analyze, and calibrate disparate sources of information. The broad geographic, year-round coverage of eBird combined with a seamless analytical framework makes it possible to perform assessments across space and time. Similarly, without critical ancillary information describing participant search effort and information to infer the absence of species (e.g., complete checklists), we would have been unable to account for the bias of imperfect detection. For this reason, we advocate for other citizen science projects to collect ancillary information sufficient to untangle the complexities of heterogeneous observation and ecological processes.

With current data volumes, the methods presented here are best suited for broadly distributed and migratory species. These methods can be easily modified for species with smaller ranges, by modifying the AdaSTEM ensemble to have a single spatial region, or for resident species, by modifying the AdaSTEM ensemble to have a single full-year temporal season. And by modifying the spatiotemporal case-control sampling and increasing the number of base models in the ensemble, the analysis can be extended to less common species. While the focus of the analysis presented here has been on pattern discovery and description, the same ensemble framework can also be modified to support confirmatory analysis by using base models that support hypothetico-deductive analysis.

The potential to use eBird data to generate robust information on species’ distributions and abundance will grow with the increasing volume and density of the data. This will make it possible to extend both the taxonomic and geographic scope of analysis. It will also improve the precision and spatiotemporal resolution of trend estimates across a wider geographic area than is currently possible. However, these improvements will be limited by the data densities in the years in which the trends begin and controlling for biases associated with the strong inter-annual increases in data volume remains a technical challenge. The increasing availability of population trends during the non-breeding season will help to refine our understanding of where and when populations are limited or regulated, complimenting migratory connectivity information derived from individual-level tracking data.

The comprehensive nature of the distribution and abundance information generated here can be used for other novel and important applications. With complete full annual cycle information, it is straightforward to make population-wide comparisons and to coordinate conservation activities across regions and seasons. Moreover, once regions of interest have been identified, the spatial resolution these estimates can be leveraged to seamlessly compare and prioritize landscapes within regions (e.g. Reynolds *et al*. 2017). In addition, the impact of regional and seasonal scale processes can be integrated across space throughout the year, making it possible to carry out accurate multi-scale population-wide impact assessments. This is important for studying a variety of broad-scale environmental and anthropogenic effects, many of which are themselves multi-scale processes, from land-use change to ecosystem services (e.g., La Sorte *et al.* 2017). The potential of our approach to integrate effects also addresses an important multi-scale challenge in climate change studies (Ådahl *et al.* 2006; Small-Lorenz *et al.* 2013) where nearly all facets of climate (e.g. temperature and precipitation) exhibit strong regional-scale intra-annual variation.

Given sufficient data density, the approach presented here can be used to leverage the broad coverage of eBird data to generate distribution and abundance information for many other bird species in different regions of the world. We have used the methods described here to analyze over 100 North American species, representing a taxonomically diverse group of species (https://ebird.org/science/status-and-trends). These results demonstrate the ability to generate robust inferences about species’ ranges, occurrence and abundance, habitat associations, and seasonal trends.

## (e) Acknowledgements

We thank the eBird participants for their contributions, the eBird team for their support, Frank A. La Sorte and reviewers for their constructive suggestions. This work was funded by The Leon Levy Foundation, The Wolf Creek Foundation, NASA (NNH12ZDA001N-ECOF), and the National Science Foundation (ABI sustaining: DBI-1356308; computing support from CNS-1059284 and CCF-1522054) and supported by the AWS Cloud Credits for Research program.

## Supporting Information

Supporting information is presented on the following topics:

1. Ensemble design,
2. Spatiotemporal Sampling procedures,
3. Base model boosted regression tree parameters,
4. Subsampling procedures to estimate uncertainty of the occurrence and abundance estimates,
5. Estimating Area of Occurrence,
6. Local Trend Estimates, and
7. Trend Simulation Model and Study Design.

Each of these sections is self-contained, with its own Literature Cited and Figures.

### Section S1: Ensemble Design

The ensemble of stixels was designed as a Monte Carlo sample of 100 randomly located spatiotemporal partitions of the spatiotemporal study extent. This resulted in a sample of stixels distributed throughout the study area that are adapted in size based on data density. Each of the 100 grids were spatially randomized by jittering the latitude and longitude grid origins and randomly rotating the resulting grid. Each of the grids were temporally randomized by jittering the starting date of the temporal grid division. Each location in space and time therefore is a member of 100 different stixels, jittered in space and time. Averaging across this sample helps control for biases associated with the arbitrary partitioning of data into stixels.

The number of stixels used to compute a local estimate across the ensemble is called the *ensemble support*. Ensemble support is important because it determines effectiveness of ensemble averaging to control inter-model variability. In this application, the maximum ensemble support is 100, the number of randomized partitions. We selected this value based on both statistical considerations and out computational budget. When training sample sizes are too small to fit base models, those stixels are dropped from the ensemble and the number of base model estimates available for ensemble averaging also decreases, increasing the variance of the ensemble estimator. We required an ensemble support of at least 75 stixels, or base models, to generate the weekly estimates of occurrence and abundance. In general, ensemble support follows patterns of data density, filtered through a combination of the base model minimum sample size requirements and stixel geometry.

Stixel size controls an important bias-variance tradeoff (Fink *et al* 2010; Fink *et al.* 2013). Stixel size needs to be chosen small enough to capture local predictor-response (i.e. species-environment) relationships, controlling the bias of base model estimates. Stixel size also needs to be chosen large enough to meet the minimum sample size requirements necessary for fitting the base models: this controls the variance when averaging across the ensemble. We began by specifying the temporal dimension of the partitions to be 366 days divided by 12 temporal partitions, equaling 30.5 contiguous days, the start days of which are then randomly jittered. An approximately 30 day window is small enough to capture a wide variety of migration patterns across a diverse set of terrestrial species using eBird data (Johnston *et al.* 2015; La Sorte *et al.* 2017). The spatial dimensions were adaptively sized to generate smaller stixels in regions with higher data density using QuadTrees (Samet 1984), a recursive partitioning algorithm. In QuadTrees, the splitting rule controls the recursion. The splitting rule was set to recursively split stixels with more than 15,000 checklists. Given high variation in data density, this splitting rule generated overly large stixels in data poor regions and extremely small stixels in areas of high data density. To prevent this, we constrained the partitioning to 1) not split stixels smaller than 5° longitude x 5° latitude regardless of the number of checklists within the stixel, and 2) forced stixels larger than 25° longitude x 25° latitude to split regardless of data density within the stixel. Fig. S1 shows the locations of eBird checklists across the study extent and Fig. S2 shows realizations of two randomly located adaptive spatial partitions used to define AdaSTEM stixels for the analysis.

**Figure S1:**
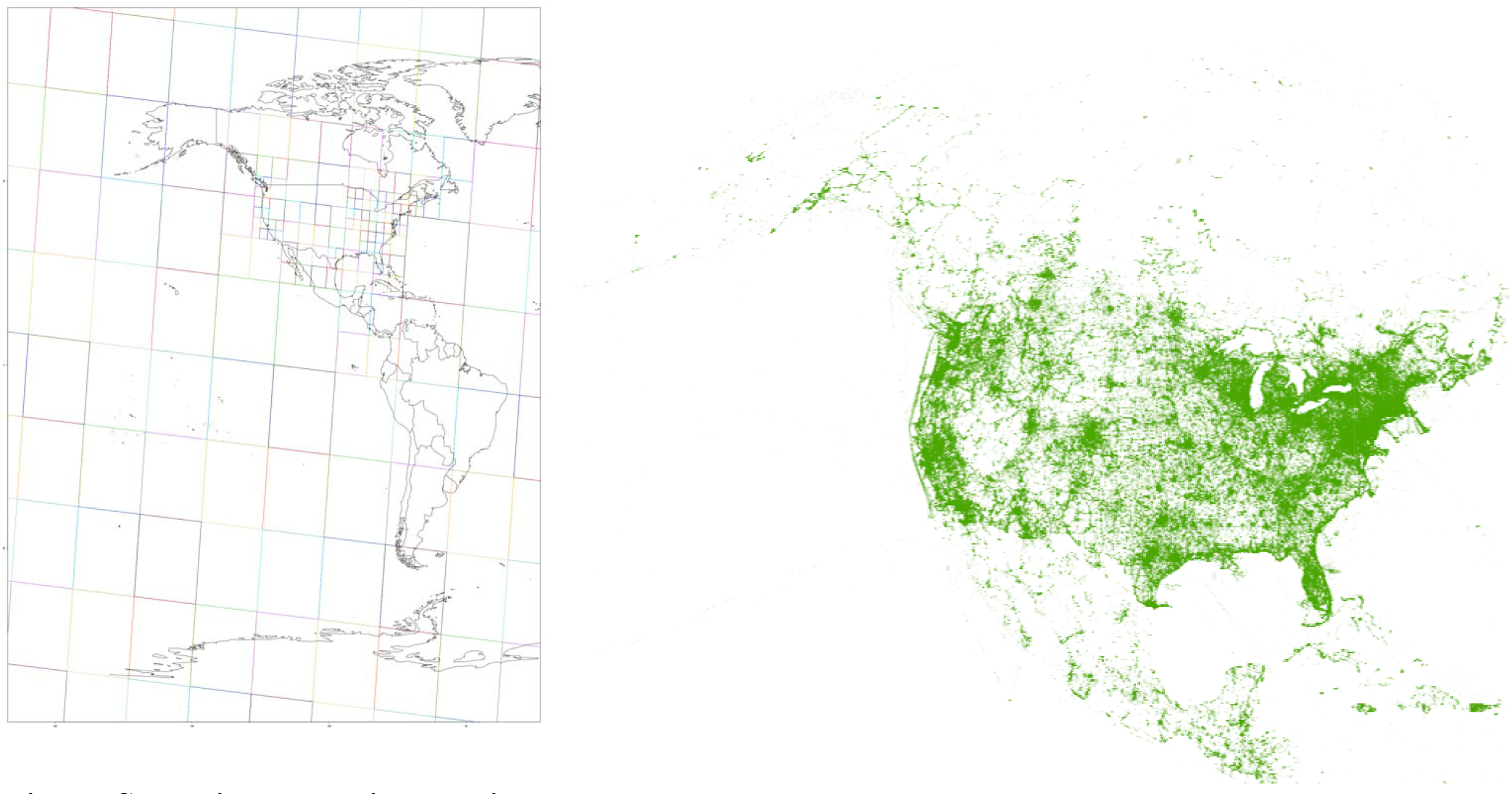
eBird checklist locations between January 1, 2004 to December 31, 2016. The spatial density of the 11.7 million checklists used in the analysis vary significantly across the North American continent.

**Figure S2:**
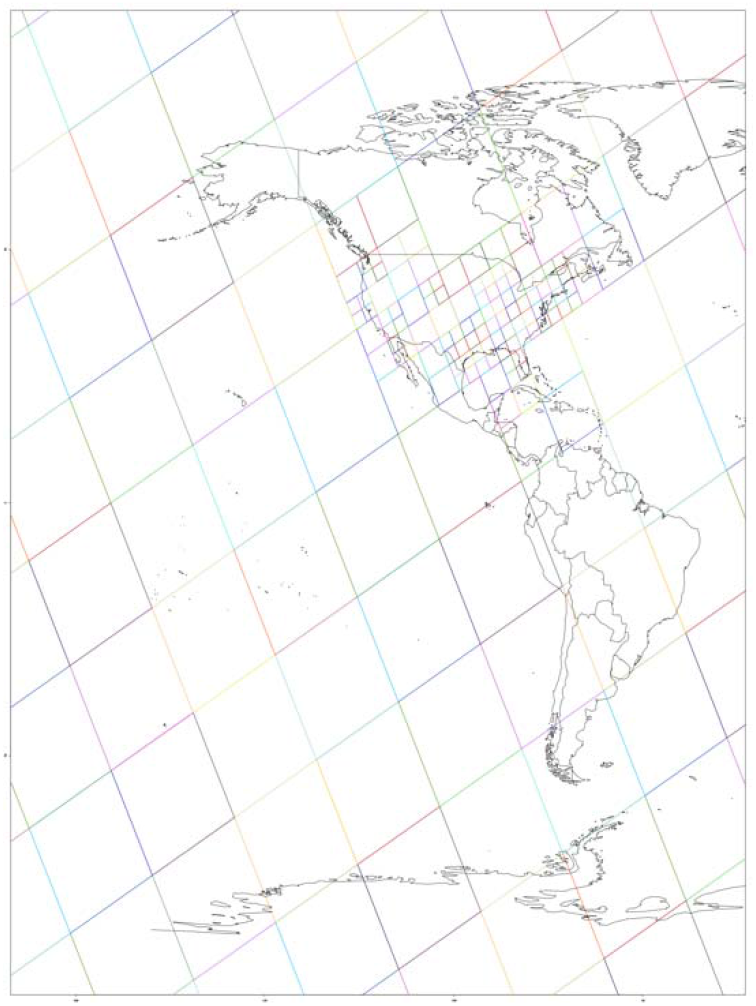
Two realizations of randomly located adaptive spatial partitions used to define AdaSTEM stixels. These image shows how stixel size adapts to data density and how stixel location is randomized through jittering and rotation. One hundred randomized adaptive partitions were used for the analysis. For each spatial partition, the temporal partition is fixed to 30.5 days, but has a randomized starting date.

## Section S2: Spatiotemporal Sampling

Within each stixel, a spatial case-control sampling strategy was used to address the challenges of highly imbalanced data and site selection bias. Imbalanced data arise when there are a very small number of species detections and a very large number of non-detections. This is a modeling concern because binary regression methods, like the first component of the ZI-BRT model, become overwhelmed by the non-detections and perform poorly (King & Zeng 2001, Robinson *et al.* 2017). The low detection rates of many species, especially along range boundaries, can result in highly imbalanced training data. This makes data imbalance a defining challenge for broad-scale, year-round modeling. By sampling detection and non-detection cases separately, case-control sampling (e.g. Breslow 1996, Fithian & Hastie 2014) improves data balance and model performance. Additionally, to alleviate spatial biases caused by the eBird site selection process, spatiotemporally balanced samples were drawn as part of the case-control sampling.

To generate spatially and temporally balanced samples for the case-control sampling, data were drawn from a randomly located regular grid, with one checklist randomly selected per 10km × 10km x 1week grid cell, applied separately for each year and separately for detection and non-detection cases. The 10km spatial grid dimension was selected to reduce the impact of repeated checklists from popular sites.

Additionally, detection data were over-sampled, using the same spatiotemporally balanced procedure, when they represented less than 25% of the balanced data. Oversampling generates ties, or repeated observations, among the training data. Because boosting, used in the ZI-BRT base models, is driven more by the set of distinct data data points than the number of tied data points generated, Mease et al. (2007) suggested breaking ties to force the boosting algorithm to respond to oversampled data. To do this we mimic the effects of imprecisely recorded checklist locations, and jitter all the spatial covariate values for each of the replicated oversampled checklists. Finally, because over-sampling detections changes the training data prevalence we correct for this change when predicting occurrence (King & Zeng 2001).

For the trend base models, we also balanced the per year sample size, after spatiotemporal case control sampling, to control for the strong inter-annual increases in eBird data volume, 20-30% per year since 2005. First, we computed the average sample size per year from 2007-16, the ten-year period over which trends were estimated. Years with less than the average sample sizes, were over-sampled (i.e. randomly sampled with replacement) and years with more than the average sample size were under-sampled (i.e. randomly sampled without replacement). This sampling strategy, what we call a reverse-mullet, resulted in a training data set with the same per-year sample size.

## Section S3: Base Model Boosted Regression Tree Parameters

Within each spatiotemporal block, we fit a two-step boosted regression tree model designed to deal with zero-inflation to predict the observed counts (abundance) of each species. The boosted regression trees for both steps of the zero-inflation model were fit with the gbm package.

The strategy used to select the base-model parameters was based on statistical considerations under the constraint of a fixed computational budget. By relying on the variance-reducing properties from averaging across the ensemble, we did not need to worry about overfitting individual base models and could avoid costly base model cross validation to select gbm parameters. This facilitated a strategy geared towards learning as much of the signal as possible with a limited number of gbm trees (ntrees = 1000) for each base model. Based on experimentation fitting base models across a set of regions, seasons, and species we set bag fraction = 0.80 and learning rate or shrinkage = 0.05. The tree.depth parameter was set to 5 for the occurrence model and 10 for the abundance model, giving both models the ability to adapt to nonlinear and interacting predictor effects.

## Section S4: Subsampling procedures to estimate uncertainty of the occurrence and abundance estimates

We used the upper 5% trimmed mean to estimate the expected occurrence and abundance across the ensemble because it is a robust estimator that guards against positive bias. A straightforward, brute force approach to estimate the uncertainty for the ensemble mean can be computed by bootstrapping the ensemble trimmed means. However, because fitting the ensemble already entails fitting 100 base models, this approach is computationally prohibitive. Instead, we employed a subsampling approach (Politis *et al.* 2009), creating ensemble replicatese by subsampling the base models.

We faced two challenges implementing this approach. First, the sample size, here, the ensemble support, was realtively small, 75–100. Second, the computational efficiency of the approach was very important because we needed to compute uncertainty estimates for up to 676M quantities per species (6.5M locations * 52 weeks * 2 estimates per location [1 for both occurrence & abundance] + 14M training & testing checklists * 2 estimates per checklist [1 for both occurrence & abundance]). To deal with these challenges we followed the computational strategy of Geyer (2013) and selected a set of parameter settings that balanced the quality of the interval estimates with the computational costs of generating them. We computed estimates of the upper 90^th^ lower 10^th^ confidence limits by subsampling with two different sizes and then computing a rate parameter correction to adjust for the original ensemble support. Following Geyer (2013), we subsampled with the square root of the ensemble support and the −1.5 power of the ensemble support.

To check these parameter settings, a small simulation test was run. We found that for sample sizes of 25 or less, the rate parameter estimates tended to be too small, resulting in intervals that were too small and had poor coverage. To mitigate this, we adjusted the rate parameter estimate upwards by 0.5 of the rate parameter’s standard error, producing more conservative uncertainty estimates. In the cases where the rate parameter estimate was negative, subsampling was not performed and quantiles of the entire sample were used producing conservative uncertainty estimates. Note that ensemble support requirements for the occurrence and abundance estimates, between 75 and 100, excludes most of these small sample size complications.

## Section S5: Estimating Area of Occurrence

At the base model level, a location was considered to be occupied if the predicted occurrence probability was above the kappa-maximized threshold for that base model. Aggregating across the ensemble, a location was considered to be occupied if at least 1/7 of base models predicted it was occupied. This is equivalent to an expert observer detecting the species at least once during 7 adjacent days of standardized surveys (estimated probability of detecting species per survey > 1/7), taking account of the variation across base models. The un/occupied status was estimated for all weeks using both the 2.8km and 25.2km spatial grids, described above.

Formally, let *m*_*s,k*_ be the estimated occurrence rate at spatiotemporal location *s* from base model *k*, *k* = 1,…*N*_*s*_, and *Kappa*_*k*_ be the kappa-maximized occurrence threshold for base model *k*, where *N*_*s*_ is the ensemble support, the number of overlapping base model estimates of location *s*. The indicator

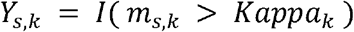

is used to estimate occurrence for location *s* for base model *k*. The sum measures how frequently the site was estimated to be occupied, 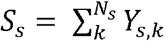. Using the fact that overlapping base models are trained using independent subsamples, we model *S*_*s*_ ~ *Binomial*(*N*_*s*_, *μ*) where *μ* is the probability of detecting at least one individual of the species on a standardized survey. To infer if the location was occupied we conducted a binomial test H0: *μ* < 1/7 This testing value was motivated by the practical consideration that a site should be considered to be occupied by a species if an expert observer can detect the species at least once during 7 adjacent days of standardized surveys.

To estimate the AOO across each species’ range for a given week often requires conducting these tests across a large number of locations. In order to identify as many occupied locations as possible while still maintaining a low false positive rate, we used False Discovery Rate (Benjamini & Hochberg 1995) thresholding to control for multiple comparisons. The q-value was set 0.01, limiting the expected proportion of falsely occupied locations to 1%. One benefit of this ensemble estimator of AOO is that it naturally adapts to regional and seasonal variation in species prevalence and detectability.

## Section S6: Estimating Local Trends

To estimate the average annual rate of change in a species’ relative abundance with moderately high spatial resolution (25.2km x 25.2km) we used a two-step approach that exploits the ensemble structure of AdaSTEM. In the first step, hypothesis tests utilize the variation across the ensemble to filter out regions where base model estimates of the trend direction were inconsistent. We call this step the *signal filter*. Then in the areas that passed the signal filter, we averaged across the ensemble to estimate the average per year change in population size while removing the intra-ensemble variation.

In the first step of the signal filter, the direction of the slope for the log-linear regression of relative abundance on year are estimated for all locations in all base models. Then, at each location, a hypothesis test is conducted to determine if the trend is increasing or decreasing. Formally, let *β*_*s,k*_ be the slope estimated at spatiotemporal location *s*, a single location in the 25km spatial grid, for base model *k*, *k* = 1,…*N*_*s*_, where *N*_*s*_ is the ensemble support, the number of overlapping base model estimates at location *s*. The slope parameter *β*_*s,k*_ describes the average annual rate of change as the percent change per year in relative abundance. To test the direction of the slope, we use a very similar approach to that used to estimate AOO (See Appendix S5). Let *D*_*s*_ = ∑ *d*_*s,k*_ where *d*_*s,k*_ = *I*(*β*_*s,k*_ > 0) indicates the direction of the trend. We model *D*_*s*_ ~*Binomial*(*N*_*s*_, *δ*_*s*_) where *δ*_*s*_ is the probability of an increasing trend at location s. When the trend is consistently estimated to be increasing across base models, *δ*_*s*_will have values close to 1, and when the trend is consistently estimated to be decreasing it will have values close to 0. To infer when trends are consistently increasing and decreasing we conduct both one-sided binomial tests H0: *δ*_*s*_ < 0.5 and H0: *δ*_*s*_ > 0.5. Because species ranges often encompass many locations across the 25km grid, ge number of tests may be conducted. In order to identify as many locations with non-zero trends as possible while still maintaining a low false positive rate, we used FDR with *q* = 0.01

Finally, we computed the ensemble averaged estimate of the trend. Like the first step, the trend was estimated as the slope here *τ*_*s*_, at location *s*, from the log-linear regression of the relative abundances on years 2007-2016. Unlike the first step, the regression was fit using the ensemble average estimates of relative abundance, as described in the Estimating Occurrence and Relative Abundance Section.

We performed a subsampling analysis to assess the uncertainty associated with sampling variation across all the steps involved in estimating local trends. Because each of the base models was trained independently using independently subsampled data sets, sampling variation of the relative abundance estimates is captured among the base model estimates. Thus, to assess the uncertainty of the trend estimates, we computed 500 replicate ensemble trend estimates, each based on a random sample of 25 out of the 100 available base models. Unlike the subsampling procedure used to estimate uncertainty for the occurrence and abundance estimates (Appendix S4), we chose *not* apply any sample size corrections to produce conservative estimates of uncertainty.

**Figure S1:**
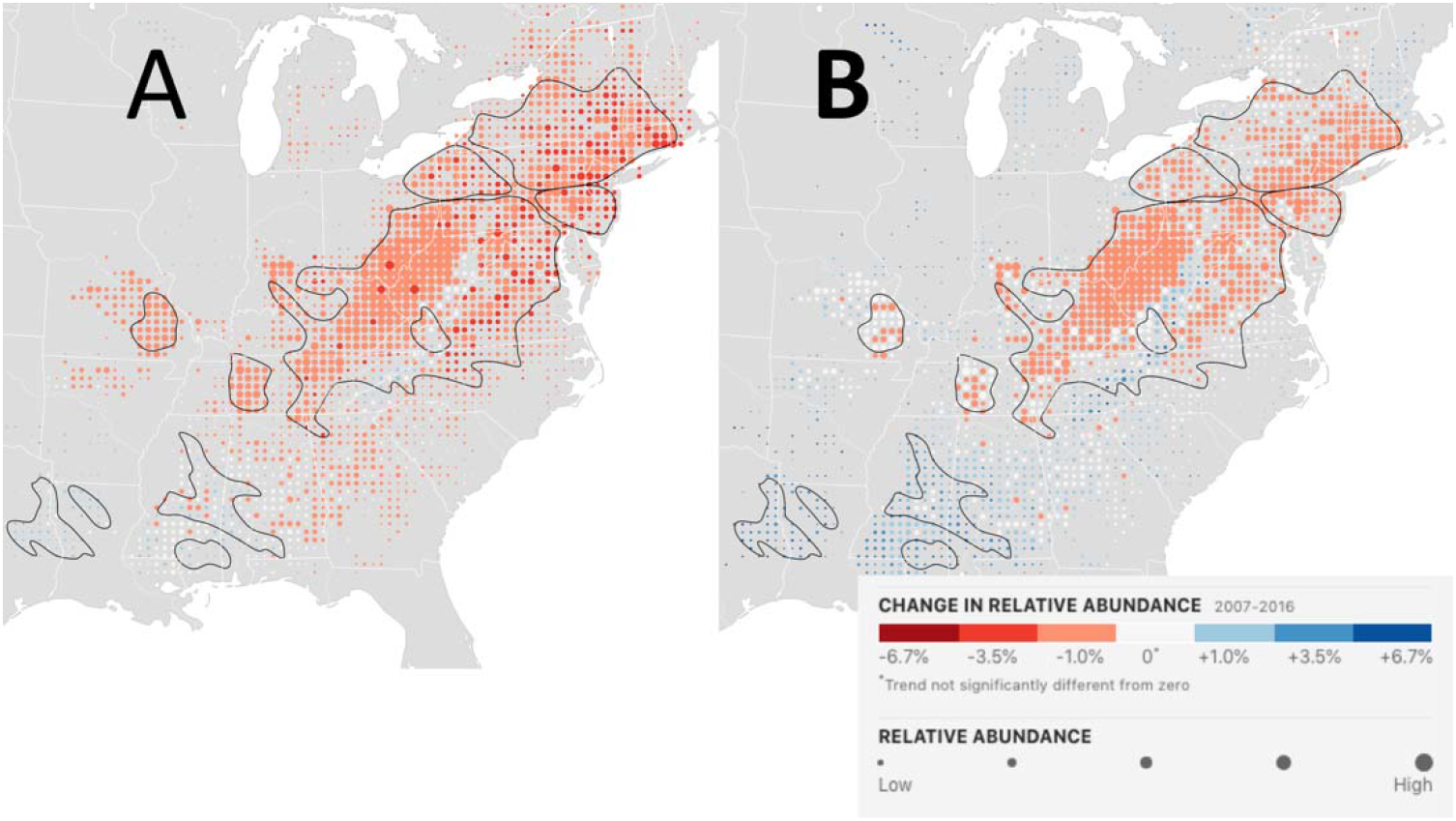
Wood Thrush 97.5% upper and 2.5% lower bound breeding season trend maps. The maps show the location-wise (A) 97.5% upper and (B) 2.5% lower limits for the average percent change per year in relative abundance from 2007–16 during the breeding season (May 30– July 3). Each dot on the map represents a 25km x 25km area. To help visualize the relative change in population size at each location, the size of each dot has been scaled according to the average abundance at that location during the 10-year study period. The black contour lines delineate those regions across which the expected False Discovery Rate is at most 5% when identifying locations with increasing and decreasing trends.

**Figure S2:**
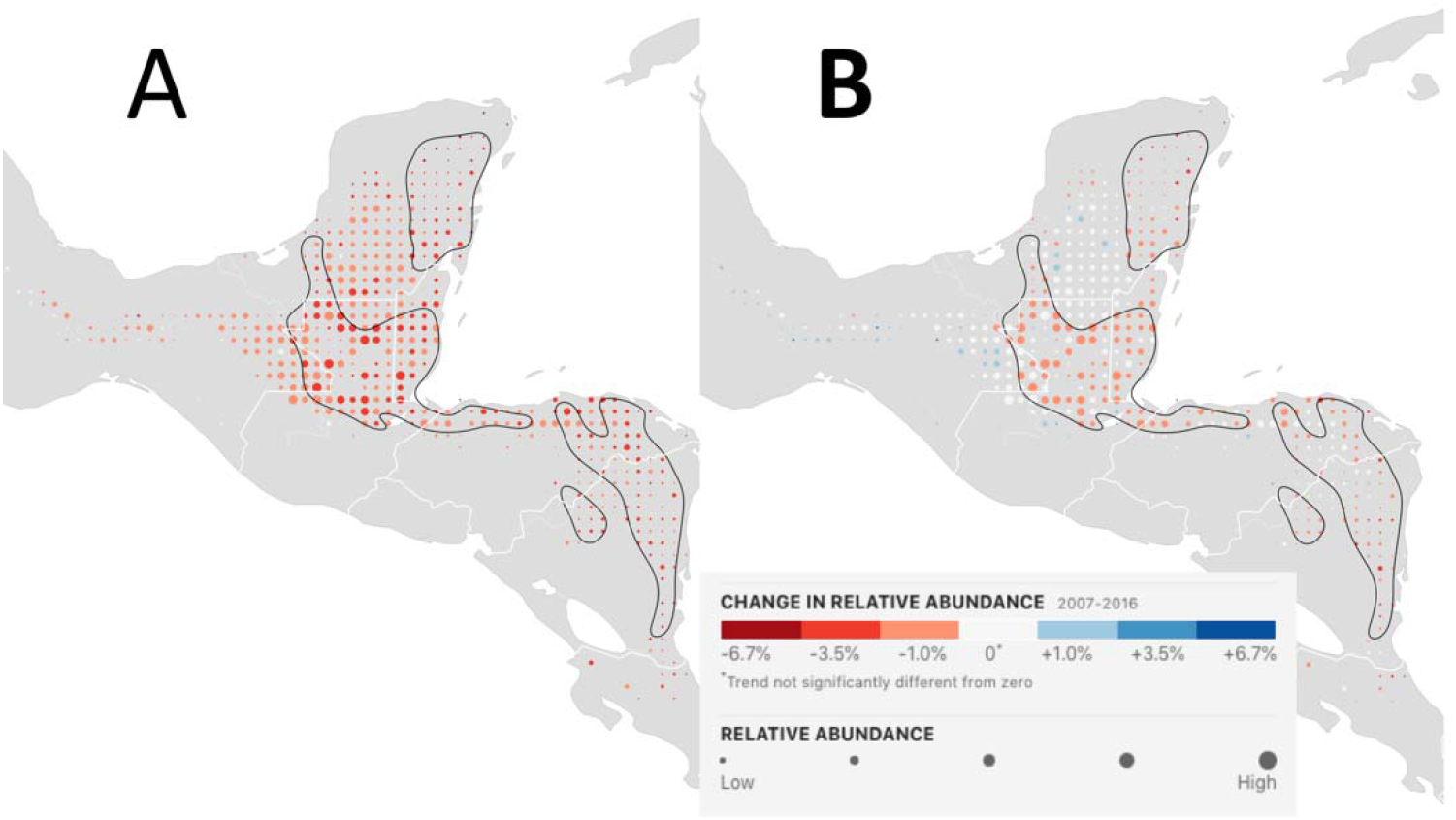
Wood Thrush 97.5% upper and 2.5% lower bound nonbreeding season trend maps. The maps show the location-wise (A) 97.5% upper and (B) 2.5% lower limits for the average percent change per year in relative abundance from 2007–16 during the breeding season (Dec 1–Feb 28). Each dot on the map represents a 25km x 25km area. To help visualize the relative change in population size at each location, the size of each dot has been scaled according to the average abundance at that location during the 10-year study period. The black contour lines delineate those regions across which the expected False Discovery Rate is at most 5% when identifying locations with increasing and decreasing trends.

## Appendix S7: Trend Simulation Model and Study Design

A simulation study was used to assess the quality of the trend estimates over the ten-year study period, 2007-2016. The study used spatially explicit simulations to generate data with specified trends while also capturing important aspects of the species’ habitat use and the citizen science observation process, both learned from training data. The power, error rate, and bias of the signal filter were assessed along with errors between known and estimated trends. There are three steps in the study:

1. Simulate data derived from populations with known trends
2. Using simulated data, estimate trends
3. Compare known and estimated trends and record statistics to describe errors, power, and bias of estimates.

The remainder of this section describes the simulation model, how the model was used to generate simulated data, the study design, and an evaluation of the breeding and nonbreeding trend estimates for Wood Thrush.

## S1.1 The Simulation Model

The simulation model was based on a ZI-BRT as described above, modified to learn specified trends along with ecological and observational patterns in the training data. Let (*N, Y, X*_*e*_, *X*_*o*_, *year*) be the set of training data for a given region, season, and species where:

- *N* is the *n x* 1 vector of observed counts on the n surveys in the training data,
- *Y* is the *n x* 1 vector that indicates the checklists with count greater than zero,
- *X*_*e*_ is the *n x k* matrix of k predictors that describe the ecological process,
- *X*_*o*_ is the *n x j* matrix of j predictors that describe the observation process, and
- *year* is the *n x* 1 vector of the year each survey was conducted.

For the given species, the region and season selected for the trend analysis and simulation must be large enough to achieve sufficient sample sizes for good model performance, controlling variance, and small enough to assume stationarity, controlling bias. We conduct two seasonal analyses for Wood Thrush, one across the breeding range from May 30–July 3 and the second across the non-breeding range from Dec 1–Feb 28.

First, we set notation and describe the standard unmodified ZI-BRT and then we explain the modifications used for the simulation. In the first step of the unmodified ZI-BRT a Bernoulli response BRT is trained to predict the probability of occurrence:

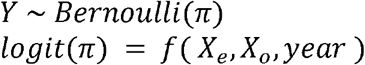

where π is the probability of occurrence and the function *f* () is fit using boosted decision trees. In the second step, the Poisson response BRT,

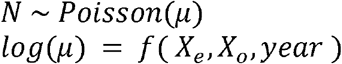

is trained to predict the expected counts *μ*, using the subset of the training data observed or predicted to be occupied.

There are some modifications to this standard ZI-BRT in order to produce the simulated data. The first modification permutes the *year* predictor variable. This ensures that the ZI-BRT cannot learn year-to-year variation from the training data and effectively removes all trends in ecology or observation process over time. The only temporal trend maintained is the increase in volume of data in later years. The second modification trains the ZI-BRT using the year-permuted training data along with an offset constructed with the specified trend. In general terms, the trend offset is *O* = *g*(*year*^*p*^) where *g* () is a function of the permuted year value, *year*^*p*^. The modified fitting procedure begins with the Bernoulli response BRT,

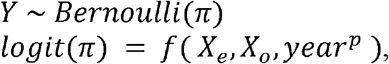

and for the Poisson response BRT is:

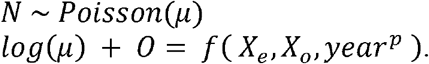

Being on the right side of this equation, the offset can be considered as an adjustment to the observed counts on the log-link scale. Thus, the boosting procedure that adaptively fits *f* () has information to estimate *g*(*year*^*p*^) from the offset.

## S1.2 Simulating New Data

After the modified ZI-BRT is trained, new data are simulated in three steps. First a new set of eBird observations is generated by sampling checklists with replacement, without regard to the search year, from the training data. Sampling this way replicates the variation observed among participant site selection, search effort, and observer effects. Year-to-year increases in the sample sizes were replicated by repeating this sampling process, independently for each year. In the second step the modified ZI-BRT is used to predict the expected occurrence and abundance, *π*\* and *μ*\*, for the set of new observations, (*X*_*0*_, *X*_*e*_, *year*)*, where the * denotes the simulated data. Finally, the binary occurrence is simulated *Y*\* ~*Bernoulli*(*π*\*) and the count, conditional on *Y*\* is simulated *N*\* ~*Poisson*(*μ*\*), generating the simulated data set, (*N*, *Y*, *X*_*e*_, *X*_*o*_, *year*)*.

## S1.3 Simulation Study Design

The simulation study was used to assess the power to detect changes in seasonal population sizes at moderately fine (25.2km x 25.2km) spatial resolution using citizen science data. Qualitatively, we want to understand how performance varies with the strength of the trend and if the method can detect spatial patterns in local trends.

To test how power varied with trend strength, simulations were constructed with increasing and decreasing trends across a range of magnitudes. To test if the method could detect spatial patterns in local trends both spatially constant and spatially varying trends were constructed. Spatially varying trends were constructed so that trend direction and magnitude varied as a function of local population density, giving rise to different trend directions at the core and edges of population distributions. Flat population trends were also included in the design to assess false positive rates. All together the study consisted of 22 combinations of spatial pattern and magnitude.

The three types of spatial trend offsets constructed were: 1) spatially constant trends, 2) spatially varying trends and 3) no trend. We used the following linear model to construct the trend offsets, *O* = *α year* + *α*_*I*_*year X*_*I*_, where α controls the strength and direction of the overall year-to-year changes in the expected log count and *α*_*I*_ controls the strength of the interaction between *year* and *X*_*I*_, the interacting variable. Note that because an intercept is fit as part of *f*(), we do not include an additional intercept term in the offset.

Spatially uniform trends were generated by setting *α*_*I*_ = 0 Trends that affect a population uniformly over a region may indicate the indirect effects of broad-spatial scale processes like climate change. Spatially varying trends can be generated by setting *α* = 0 and specifying a spatially patterned variable *X*_*I*_ to interact with *year*. To assess if spatial patterns associated with density dependent population processes can be detected, we selected *X*_*I*_ to be the PLAND cover class predictor with the largest Spearman rank correlation between itself and *π*\*, used here as an index of population density. Processes like habitat loss, disease, and dispersal can interact with population density to generate spatially varying trend patterns, e.g. Channell and Lomolino (2000) and Massimino, et al. (2015).

Using two parameter sweeps, the spatially constant models were generated with the *α* ranging from −0.08 to 0.08 in 11 values spaced 0.016 apart and the spatially varying models were generated with *α*_*I*_ ranging from −0.40 to 0.40 in 11 values spaced 0.08 apart for a total of 22 simulation *treatments*. The strongest trends were parameterized to generate relatively large regions within the species’ range experiencing changes in population size of at least 6.7% per year over 10 years, one of the IUCN red-list criteria for endangered populations (IUCN 2019).

## S1.4 Simulation Evaluations

Trend estimation proceeds in two steps as described above, where the signal filter first detects local trends and the trend magnitude is estimated in locations where the direction of trends is consistent. For each simulation we evaluated the power, error rate, and bias of the signal filter along with the correspondence between the magnitude of known and estimated trends. The false detection proportion (FDP) was calculated as the number of locations on the 25km grid where trends were erroneously detected, as a proportion of the total number of locations where trends were detected. The power was calculated as the proportion of locations where a trend was correctly identified out of all locations known to have non-zero trends. To understand how power varied as a function of the local trend strength, power was also evaluated across all locations with known trends with a minimum magnitude, ranging from 0 to 15% per year. Where the signal filter detected local trends, the coefficient of determination (*R*^2^) was computed to describe the proportion of variation in the known magnitudes explained by the estimates.

For each of the breeding and non-breeding seasons, a separate simulation study was conducted for each of the 22 simulation treatments. For each treatment, the training data was spatiotemporally sampled and the year predictor variable was permuted to fit the simulation model. The AdaSTEM base models for each simulation treatment were trained using 100 independent realizations of simulated data.

We measured the performance of the trend estimates averaged across the full suite of simulation treatments to estimate the expected performance across a wide variety of trend scenarios. An important part of this assessment was quantifying directional biases when detecting trends. When biases were found, we adjusted the signal filter to provide robust control against false detection of trends and conservative power estimates.

If the FDP was found to exceed a specified error limit (e.g. 5, 10 or 20%) for more than 10% of the of all the locations in all of the simulations, we considered the trend estimator to be biased for that error limit and season. To quantify and adjust for this bias we modified the directional hypotheses used for the signal filter, *H*0: *δ*_*s*_ < (0.5 − *B*^−^) and H0: *δ*_*s*_ > (0.5 + *B*^+^) where parameters *B*^+^ and *B*^−^ ∈ (0,0.5] describe the directional biases. As the values of each bias parameter increases, the signal filter requires more consistency in the direction of the trend estimates across the ensemble, thereby reducing the FDP. Similarly, as the values of each bias parameter increases, the fewer locations on the trend map where trends can be identified while guaranteeing the FDR at the specified error limit.

To estimate the directional biases, we performed a parameter sweep evaluating FDP and power across all combinations of values of *B*^+^, *B*^−^ ∈ (0.0, 0.01, 0.02,…, 0.25). Then we estimated the value of *B*^+^, *B*^−^ that maximized power subject to the constraint that FDP was less than the specified limit (e.g. 5, 10, or 20%) across ≥ 90% of the simulations.

All of trend estimates reported here, for both breeding and nonbreeding seasons, were made using a FDR limit of 5%. For each of the breeding and nonbreeding simulations we estimated the direction bias parameters (*B*^+^, *B*^−^) and used them to estimate trends. Thus, all of the trend maps, power statistics, and *R*^2^ measurements reported in this paper were made using these bias corrections under a 5% FDR limit.

## S1.5 Wood Thrush Simulations

Two simulation studies were conducted for the Wood Thrush over the 2007-2016 study period, one for the breeding season (May 30–July 3) across the species’ range in the northeastern North America and the second for the non-breeding season (Dec 1–Feb 28) across the species’ range in Central America.

The simulations provide qualitative information describing the ability of the method to identify spatially varying trend patterns among locations. Fig. S1–4 show simulated and estimated trend maps for a sample of simulation treatments across a broad array of spatially constant and spatially varying trends with trend magnitudes that vary in direction and magnitude. The trend magnitudes varied along the rows of each figure with weak (includes regions with trends ~|1%/*yr*|), medium (includes regions with trends ~|3.5%/*yr*|), and strong (includes regions with trends ~|6.7%/*yr*|) trend magnitudes. This suite of spatial trend patterns is varied enough to begin to assess the method’s ability to estimate spatial patterns across locations. The quality of the trend estimates improves from weak to strong trend magnitudes, regardless of spatial pattern or direction. Regional patterns are identified, though with errors, when simulated trends are weak, and become clearer as trends become stronger. The magnitude of the estimates generally varies with simulated trend strength, visible as the correspondence between the darkness of the colors shown for the estimate and simulation trend map pairs in Fig. S1–4. However, in regions with declining trends the trend magnitude appears to be underestimated in the nonbreeding season and among the spatially varying treatments in the breeding season.

Fig. S5 shows power curves as a function of the minimum simulated trend magnitude, for 5, 10, and 20% FDR constraints for both seasonal simulation analyses. Both plots show the expected pattern of increasing power with increasing minimum trend magnitude. The plots also show the expected tradeoff between FDR and power, with increasing power as the FDR constraint becomes more lenient. We recognize that in some conservation applications the false detection of declining trends, carries a far lower risk for a species than failing to detect a declining trend, and in such circumstances, it may make sense to increase the error limit to 10 or 20% to improve the power to detect trends as can be seen in Fig. S5.

Finally, the correspondence between estimated and simulated known trend magnitudes was stronger in the breeding season (*R*^2^=75.6%) than the nonbreeding season (*R*^2^=59.5%). Overall, these simulation results suggest that breeding season trend estimates will be more accurate, powerful, and less variable than those in the nonbreeding season. In general, this is expected because of the much higher density of data across the breeding range compared to the nonbreeding range.

**Figure S1:**
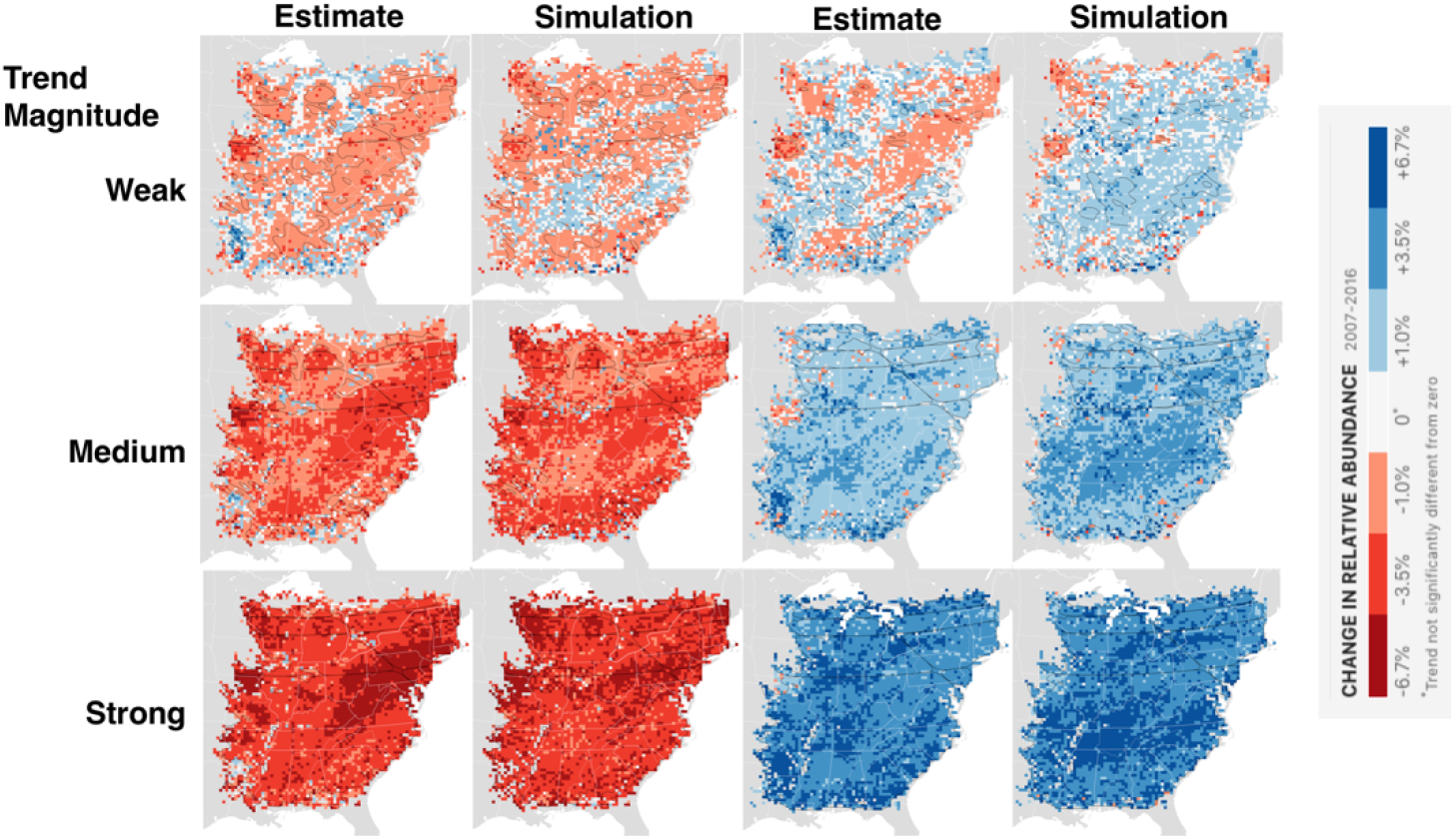
Wood Thrush breeding season simulated and estimated trend maps for spatially constant treatments. The trend magnitude varies along the rows with weak (includes regions with trends ~|1%/*yr*|), medium (includes regions with trends ~|3.5%/*yr*|), and strong (includes regions with trends ~|6.7%/*yr*|) trend magnitudes. The first two columns show estimated and simulated trends for decreasing trends. The third and fourth columns show estimated and simulated trends for decreasing trends. The black contours delineate the regions across which the expected False Discovery Rate is at most 5%.

**Figure S2:**
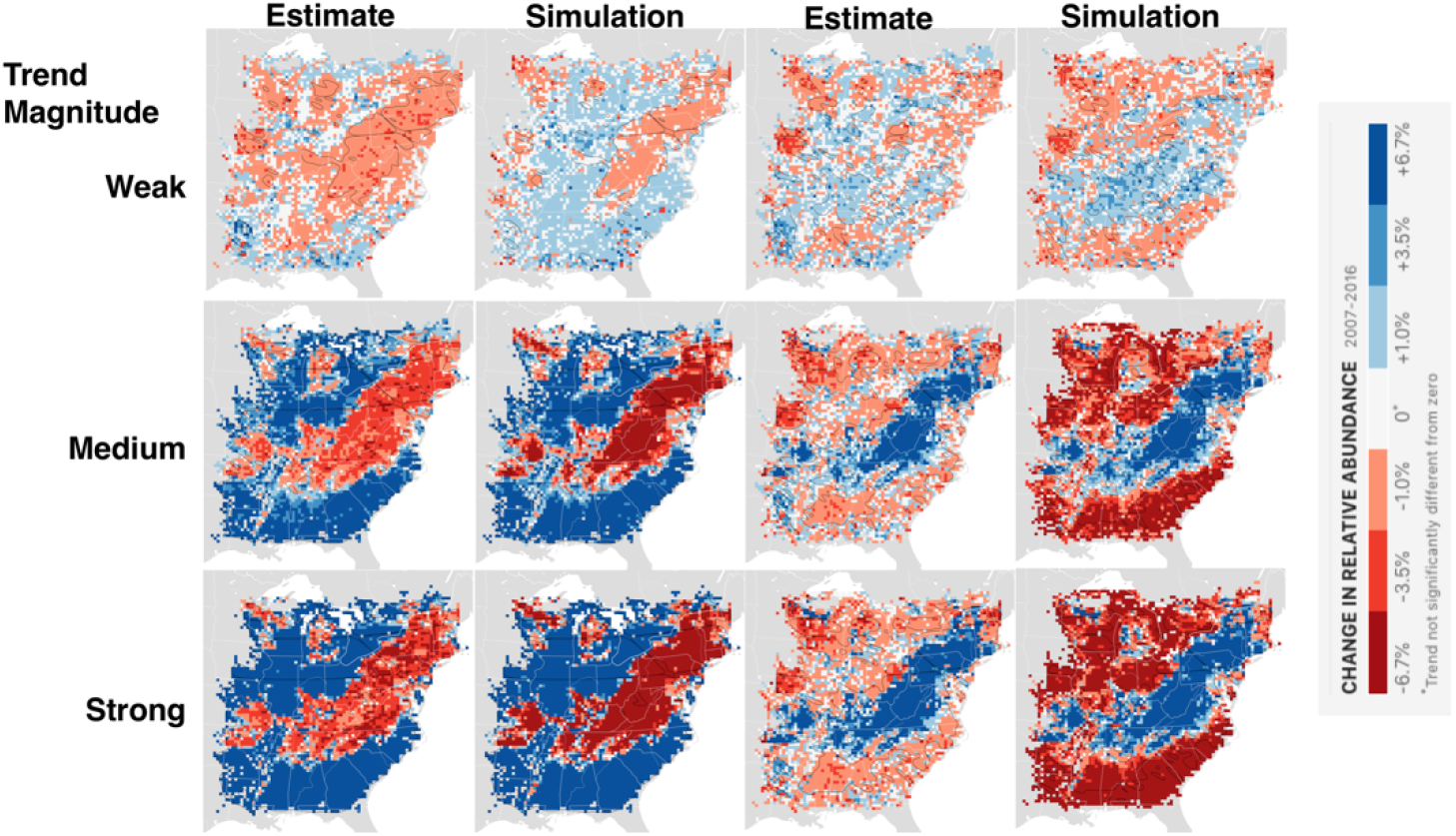
Wood Thrush breeding season simulated and estimated trend maps for spatially varying treatments. The trend magnitude varies along the rows with weak (includes regions with trends ~|1%/*yr*|), medium (includes regions with trends ~|3.5%/*yr*|), and strong (includes regions with trends ~|6.7%/*yr*|) trend magnitudes. The first two columns show estimated and simulated trends for decreasing trends. The third and fourth columns show estimated and simulated trends for decreasing trends. The black contours delineate the regions across which the expected False Discovery Rate is at most 5%.

**Figure S3:**
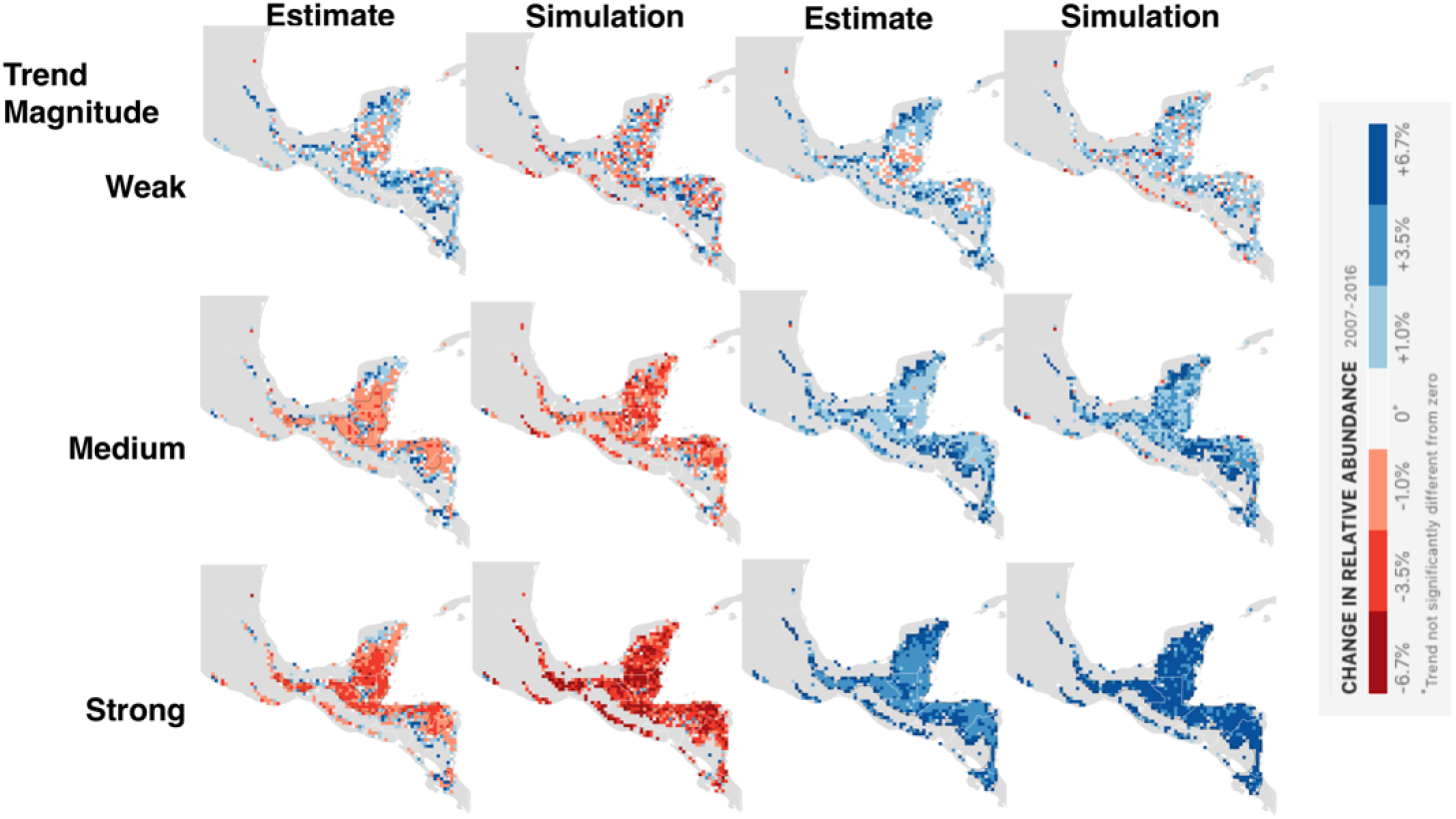
Wood Thrush nonbreeding season simulated and estimated trend maps for spatially constant treatments. The trend magnitude varies along the rows with weak (includes regions with trends ~|1%/*yr*|), medium (includes regions with trends ~|3.5%/*yr*|), and strong (includes regions with trends ~|6.7%/*yr*|) trend magnitudes. The first two columns show estimated and simulated trends for decreasing trends. The third and fourth columns show estimated and simulated trends for decreasing trends. The black contours delineate the regions across which the expected False Discovery Rate is at most 5%.

**Figure S4:**
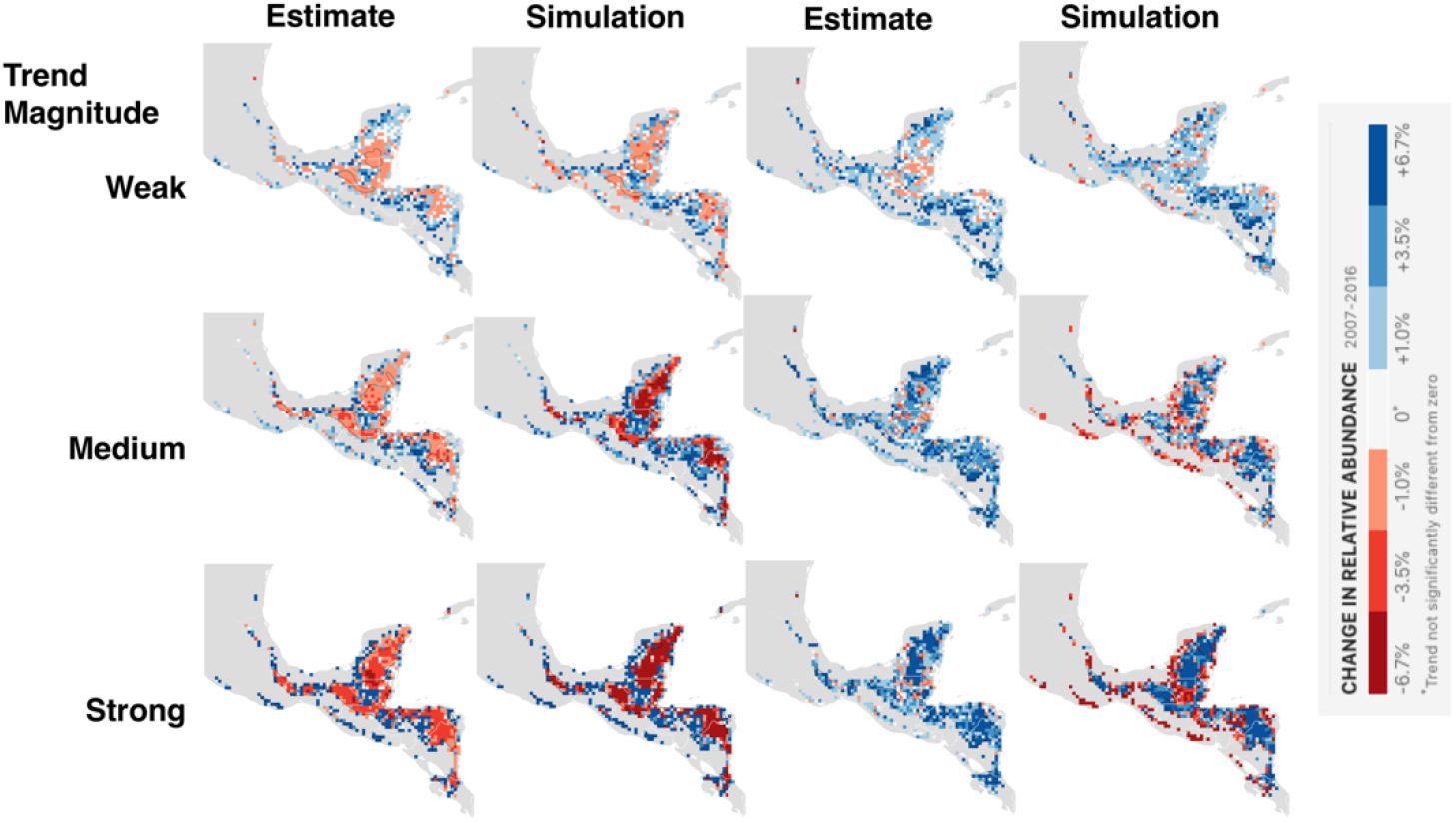
Wood Thrush nonbreeding season simulated and estimated trend maps for spatially varying treatments. The trend magnitude varies along the rows with weak (includes regions with trends ~|1%/*yr*|), medium (includes regions with trends ~|3.5%/*yr*|), and strong (includes regions with trends ~|6.7%/*yr*|) trend magnitudes. The first two columns show estimated and simulated trends for decreasing trends. The third and fourth columns show estimated and simulated trends for decreasing trends. The black contours delineate the regions across which the expected False Discovery Rate is at most 5%.

**Figure S5:**
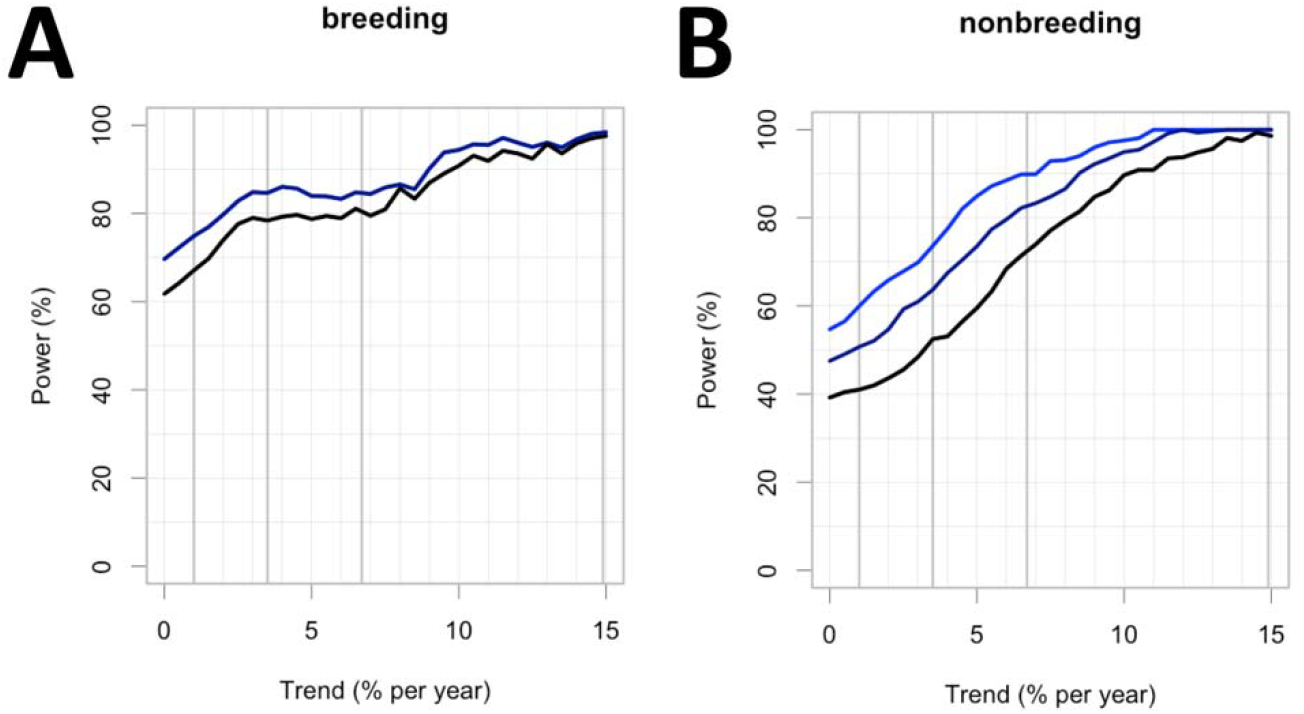
Wood Thrush seasonal power curves as a function of the minimum simulated trend magnitude. Power varies as a function of the minimum trend magnitude for the (A) breeding and (B) nonbreeding season analyses. Power is reported as the percentage of all locations in range across the simulated known map that meet the minimum magnitude requirement that were identified with the correct trend direction when FDR was constrained at 5 (black), 10 (dark blue), and 20% (light blue). The maximum false detection proportion was 8% for the breeding season, so no light blue line is shown. The overall power corresponds to a minimum trend of zero, at the leftmost side of the graph. The black line corresponds to the black contour lines in Fig. 4 and 5.

